# Flexible fitting of PROTAC concentration-response curves with changepoint Gaussian Processes

**DOI:** 10.1101/2020.11.13.379883

**Authors:** Elizaveta Semenova, Maria Luisa Guerriero, Bairu Zhang, Andreas Hock, Philip Hopcroft, Ganesh Kadamur, Avid M. Afzal, Stanley E. Lazic

## Abstract

A proteolysis targeting chimera (PROTAC) is a new technology that marks proteins for degradation in a highly specific manner. During screening, PROTAC compounds are tested in concentration-response (CR) assays to determine their potency, and parameters such as the half-maximal degradation concentration (DC_50_) are estimated from the fitted CR curves. These parameters are used to rank compounds, with lower DC_50_ values indicating greater potency. However, PROTAC data often exhibit bi-phasic and poly-phasic relationships, making standard sigmoidal CR models inappropriate. A common solution includes manual omitting of points (the so called “masking” step) allowing standard models to be used on the reduced datasets. Due to its manual and subjective nature, masking becomes a costly and non-reproducible procedure. We, therefore, used a Bayesian changepoint Gaussian Processes model that can flexibly fit both non-sigmoidal and sigmoidal CR curves without user input. Parameters, such as the DC_50_, the maximum effect *D*_max_, and the point of departure (PoD) are estimated from the fitted curves. We then rank compounds based on one or more parameters, and propagate the parameter uncertainty into the rankings, enabling us to confidently state if one compound is better than another. Hence, we used a flexible and automated procedure for PROTAC screening experiments. By minimizing subjective decisions, our approach reduces time, cost, and ensures reproducibility of the compound ranking procedure. The code and data are provided on GitHub (https://github.com/elizavetase-menova/gp_concentration_response).

## Introduction

A proteolysis targeting chimera (PROTAC) is a new drug modality that induces degradation of disease-causing proteins. It is a molecule composed of two active domains and a linker and forms complex consisting of the target, an E3 ligase, and the PROTAC molecule. PROTAC molecules ubiquitinate their target proteins, thereby tagging them for degradation by the proteasome. Factors to consider when designing PROTACs include which E3 ligase to target, the required binding affinities between the target protein and E3 ligase, the target protein basal turnover rate, and the E3 ligase expression level in the relevant tissue. Small molecules typically inhibit protein activity, but PROTACs completely remove the protein from cells. As the PROTAC molecule itself is not consumed during proteasomal degradation, it can be recycled and used many times. Catalytic knockdown of proteins in vivo has been shown in some studies^1^. Precisely estimating parameters of CR curves, such as potency and efficacy is essential for robust drug development.

Traditional small molecule screening for inhibitors typically yields sigmoidal monotonic concentration-response curves, characterized by a plateau at high drug concentrations. In contrast, PROTAC CR curves can have a distinct profile, marked by a loss of efficacy at higher doses. Accounting for such a “hook effect” in data analysis is essential to understand the underlying mechanism. A mathematical framework has been developed to understand the ternary complex equilibria^2^: it explains that the concentration-response behavior of such a complex should form a hook. The theoretical model predicts a symmetric curve with respect to the point of the maximal effect. However, experimental data frequently displays non-symmetric patterns. A common solution includes manually omitting points (the so called “masking” step) allowing standard models to be used on the reduced datasets. This approach has several drawbacks. Due to its manual and subjective nature, it is time consuming and may vary from scientist to scientist. Furthermore, even the same scientist at different time points might judge differently (due to their increasing expertise or more random factors). As a result, manual masking becomes a costly and non-reproducible procedure. There is therefore an unmet need for an automated method to fit models to experimental data allowing for non-sigmoidal shapes. Such a method should minimize subjectivity, increase reproducibility, as well as provide uncertainty quantification to the ranking of compounds.

The challenge of non-sigmoidal fits can be addressed in several ways - either by adjusting parametric models, or by non-parametric modelling. Parametric models are widely used in drug discovery as a tool to fit CR curves. They have a pre-defined functional form, such as a linear, exponential decay^3^, or Hill model^4^. The parameters governing these models often have a straightforward interpretation, such as the slope and intercept for a linear model, or the concentration that gives the half-maximal degradation (DC_50_) and the maximal effect (D_max_) for a sigmoidal model. Parametric models reduce an unknown and potentially complicated true function, to a simple form with few parameters. Such models can produce consistent results when observations follow a pre-defined class of shapes but may not fit the data well when a pattern exhibited by data does not fall into that class. Then the simplicity of a parametric model might lead to erroneous estimates. Another price to pay for the simplicity of parametric models is their global behavior, i.e. a change in one parameter can change the shape of the whole curve.

Non-parametric models such as LOESS (LOcally Estimated Scatterplot Smoothing), splines, and Gaussian processes (GPs) are more flexible and learn the shape of a curve from data. GPs have certain advantages over its non-parametric counterparts: the LOESS method is known to require fairly large amounts of data, while GPs can deal with small data sets, and splines can be viewed as a special case of GP regression^5^. Additionally, the parameters of most non-parametric models are not directly interpretable, while such models are usually more computationally intensive and harder to implement. GPs are able to carry more interpretable information than its non-parametric counterparts. For instance, its changepoint kernel can incorporate interpretable parameters, such as point of departure (PoD). PoD has been proposed as an alternative to DC_50_ for measuring potency of compounds. A few definitions of POD have been proposed in the literature^6^. Here we define it as the changepoint in the changepoint GP model, i.e. the concentration at which the curve changes shape from flat to flexible. DC_50_ measures half-degradation, and POD measures the concentration where degradation starts. In addition, some parts of a curve can be fixed while other modelled more flexibility. A parametric framework offers two approaches to fitting when the shape of a curve is not known in advance: model averaging^7^ and model selection. Both approaches start with a set of parametric models and then undergo two stages. At the first stage, models are fit to the data. At the second stage, models are either averaged, or one model is selected as the best. Both methods would not work well if the initial set does not contain a realistic model for the given data. Non-parametric approaches can avoid the two-stage process, while the necessary decision-influencing parameters can still be derived from the fitted curves.

Established methods for estimating parameters of a sigmoidal concentration-response curve are based on minimization of least squares or maximization of the likelihood via such optimization algorithms as gradient descent, Gauss-Newton and Marquardt–Levenberg^8^. These methods produce results in the form of point estimates of parameters. Classical statistical models can communicate uncertainty in the form of confidence intervals (CIs). However, they do not inform which values within the CIs are more or less probable as they only show the range of values. On the contrary, in Bayesian inference, parameters are described by probability distributions. Bayesian credible intervals (BCIs) can also be extracted, but each value within BCI is also supplied with information about how probable it is. Additionally, the Bayesian inference method allows to incorporate prior beliefs and take several sources of uncertainty into account. These features make the method a powerful and attractive tool to capture, model and communicate uncertainty. Uncertainty in model parameters can be incorporated and propagated into the estimates of derived quantities. Bayesian approach to dose-response curve fitting has started gaining recognition in the analysis of *in vivo* data.

The aim of this work is threefold: 1) to provide an automatic method for fitting curves of non-standard shapes to reduce subjectivity and improve reproducibility, 2) to supply parameter estimates and final ranking with uncertainty quantification, 3) to demonstrate the usefulness of a Bayesian changepoint GP model for fitting PROTAC CR curves. The changepoint GP kernel can infer a PoD. It captures the biology such as a flat (negligible) effect at concentrations up to the PoD, and more flexibility at higher concentrations to capture the behavior of interest. An advantage of a Bayesian framework is that the model can account for several sources of uncertainty and represents the parameter uncertainty in the form of a distribution, which is propagated to the uncertainty of the compound ranks. By minimizing the number of subjective decisions, our approach reduces time, cost, and ensures reproducibility of the compound ranking procedure. The process of compound ranking is summarized on Supplement Figure 1.

## Materials and Methods

### Data

The data was collected in a set of 2 experiments for 9 compounds as follows: 384 well plates (PE cell carrier ultra, PerkinElmer, 6057308) containing 12,000 cells per well in RPMI1640 medium (Sigma, R7638) supplemented with glutamine (Gibco, 25030) and 10% FCS were dosed with an Echo 555 (labcyte) at the indicated doses. Maximum effect and neutral controls DMSO were also used. After one day of incubation, cells were fixed with PFA (4% final concentration) for approx. 20 minutes, washed (3X PBS by using a BioTek plate washer) and subsequently stained with primary followed by secondary antibody for target detection and DRAQ5 (Abcam, Ab108410). Each staining step was followed by a washing protocol described above. The signal for the target antibody stain per well was measured by automated microscopy (Cellinsight, Thermo). Data were normalized by Genedata Screener software (Genedata Screener®) using controls distributed at several positions across the plate to 0% (DMSO, neutral control) and −100% (maximal effect compound). The core formula for normalization is the expression (*x*-<*cr*>)/(<*sr*>-<*cr*>) where *x* is the measured raw signal value of a well, <*cr*> is the median of the neutral controls, and <*sr*> is the median of the measured maximal effect control compound.

### Model

#### Bayesian inference

The Bayesian model formulation consists of two parts: preliminary information, expressed as prior distributions, and a likelihood. Inference is made by updating the prior distributions of the parameters according to the data via the likelihood. The likelihood reflects the assumptions about the data generating process and allows a model to be evaluated against the observed data. In the process of inference, samples of parameters are drawn from posterior distributions. These samples can be consequently used to quantify uncertainty of parameter estimates in the form of distributions, as well as to derive statistics.

The logic of the subsequent sections is as follows. First, we explain the likelihood of the model and how to account for different sources of variation in the section “Measurement model and likelihood”. After that, we give a brief introduction to the Gaussian Process class of models and build on the kernel proposed in the literature^9^ by generalizing it to a wider family of changepoint kernels in the section “changepoint Gaussian Processes”. We round up the model formulation by explaining our prior choices in the section “priors for model parameters”.

#### Measurement model and likelihood

We assume that both data for treatments and controls is available. Data for controls consists of *N*_*control*_ measurements of the response, corresponding to the same (zero) concentration. These measurements therefore can be denoted as pairs (*x*_*0*_, *y*_*control*_^*c*^), *c* = 1,..,*N*_*control*_. Treatment data is available for each concentration *x*_*i*_ (*i*=1,…,*N*) in several measurements. We denote such data as 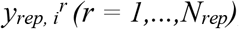, where index *i* corresponds to one of *N* concentrations, and index *r* corresponds to the replicate number. The aim of modelling is to build a curve <u>*y*</u> which captures data trend in the best way. At this point, we do not need to make any assumption about the exact functional form of <u>*y*</u>, i.e. it can be either parametric or non-parametric. Subse-quently, we call <u>*y*</u> “the mean predicted curve”. The observed points, however, will not lie on the curve <u>*y*</u> exactly. This observation can be attributed to the present sources of uncertainty. The model takes two sources of variation into account: curve uncertainty *σ* and replicate-to-replicate variation *σ*_rep_. Curve uncertainty explains why all observations do not lie exactly on the mean predicted curve, and replicate uncertainty explains why measurements at the same concentration do not coincide with each other. These sources of variation are presented graphically on Figure 1. When only one replicate is available, the treatment observations could be described as *y*_*i*_ = <u>*y*</u>_*i*_ + *ɛ*_*i*_, *ɛ*_*i*_ ~ *N*(0, *σ*^2^). Hence, it holds for *y* itself that it is distributed normally, is centered at <u>*y*</u> and has standard deviation *σ*. Using vector notation (*y* is a vector with components *y*_*i*_, *i* = 1,…*N*) we get

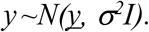

**Figure 1:**
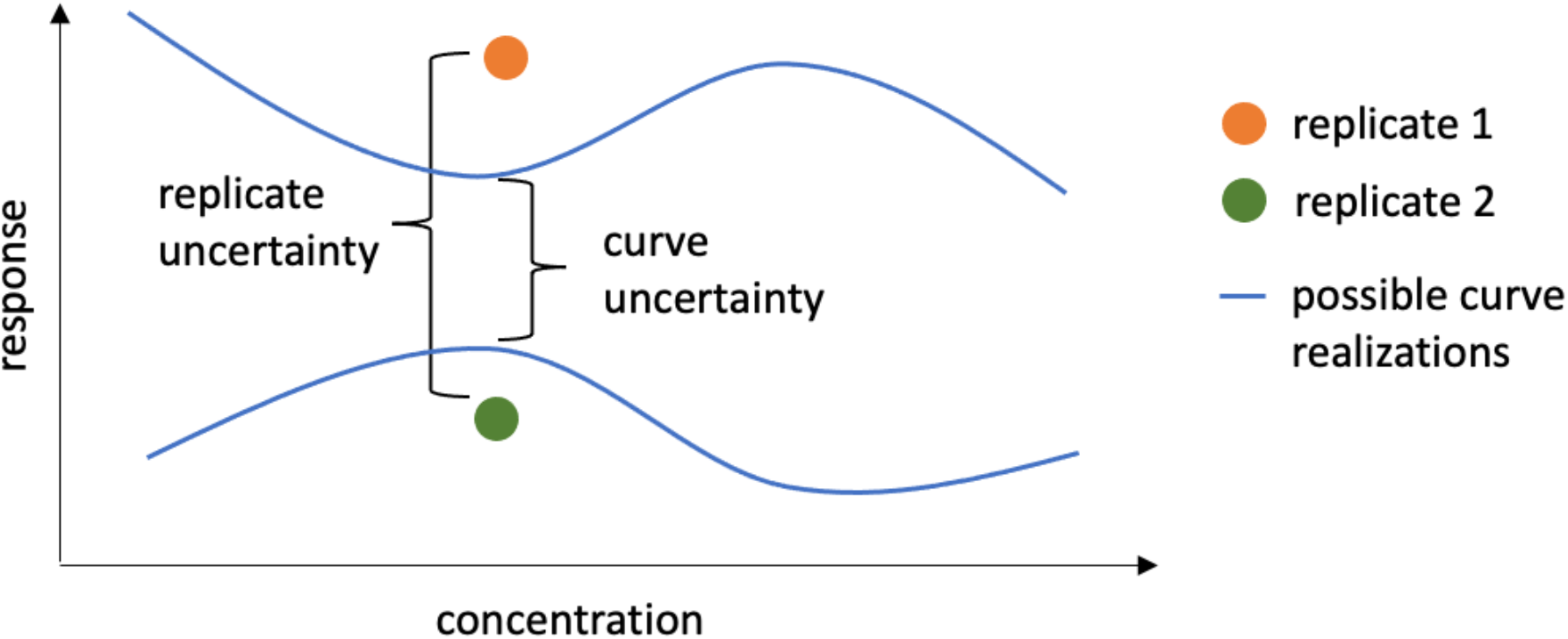
Sources of variation: curve uncertainty and replicate-to-replicate uncertainty.

Variability in replicates for the same concentration 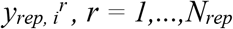 is mod-elled with a Student’s t-distribution with degrees of freedom *v*, centered on the mean response and replicate-to-replicate scale 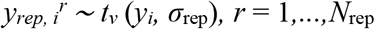. In vector notation this becomes

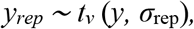

Here *v* is a parameter which gets inferred from data. Student’s t-distribution has heavier tails than the normal distribution and, hence, allows for more extreme observations. This choice makes the model less sensitive to outliers. The higher is the estimate of *v*, the closer is the distribution to normal. Hence, the estimates of this parameter can diagnose the presence or absence of outliers in the data. To summarize, treatment data is distributed as

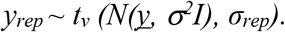

It would be hard to derive the analytical form of this distribution. However, to fit a Bayesian model the analytical form of the distribution is also not needed. What we can notice is that the total variation of the replicates approximately equals *σ*^2^ + *σ*_*rep*_^2^. Data for controls *y*_*control*_^*c*^, *c* = 1,..,.*N*_*control*_ has mean *μ* and the same two sources of variation: curve uncertainty and replicate-to-replicate uncertainty. That is why, to match the amount of modelled uncertainty in the treatments, heuristically, the likelihood for controls is derived as

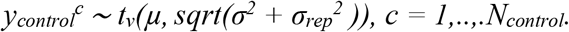

Distributions for *y*_rep_ and *y*_control_ define the likelihood of the model. The computational graph, summarizing this logic, is presented in the Supplement Figure 2.

#### Changepoint Gaussian Processes

The trend <u>*y*</u> as the dependence of the outcome variable (protein degradation) on the concentration is modelled using a GP with a changepoint kernel. A composite GP kernel has been proposed in the context of dose-response curve fitting in^9^. The proposed covariance structure is a special case of a changepoint GP. Here we present a more general framework of the changepoint kernel modelling and explain how^9^ fits into it.

A GP defines a probability distribution over functions, i.e., each sample from a GP is an entire function, indexed by a set of coordinates (concentrations). Evaluated at a set of given concentrations, a GP *f* is uniquely specified by its mean vector *μ* and covariance matrix *K* and follows a multivariate normal distribution: *f* ~ *N*(<u>*μ*, *K*</u>). Each element of the covariance matrix *K*_*i,j*_ is obtained as an evaluation of the covariance kernel – a function of two arguments *k*(. , .) at a pair of points *x*i and *x*j. The kernel controls variance-covariance structure: its value at a set of two points *k*(*x*_*i*_, *x*_*j*_) quantifies the covariance of two random variables *f*(*x*_*i*_) and *f*(*x*_*j*_). Kernel design is crucial for GP modelling since it describes the class of possible functions which can be captured: both the global shape, as well as local properties, such as smoothness^10^. Most commonly used kernels, such as exponential *η*^2^*exp*(-*d/ρ*), squared exponential *η*^2^*exp*(-*d*^2^/*ρ*^2^), or Matern *C*_*v,ρ*_(*d*), depend on the distance *d* = || *x*_i_ – *x*_j_|| between two points and, hence, are stationary. They are inappropriate for modelling concentration-response curves and surfaces (e.g., drug-drug interaction) which might display different behavior across the observed concentrations. Kernels derived from exponential and squared exponential kernels would have smooth shapes, while Matern kernels under certain values of *ν* result in curves with more local fluctuations and less smoothness. The length-scale *ρ*, common for many kernels, governs the flexibility of the curves: smaller length-scales induce correlations between points at smaller distances and hence more oscillatory behavior - such behavior, when exaggerated, can lead to overfitting; too large length-scales, on the other hand, make curve fits too inflexible and may lead to underfitting the data. The amplitude *η*, frequently applied to kernels as a constant multiplier, controls the expected variation in the output. The amplitude controls how strong the functions fluctuate and the length-scale controls how rapidly they fluctuate. Dependence of the standard kernels on their parameters can be visually inspected in^11^. Choice of priors of the GP parameters plays a big role in their identifiability.

Covariance functions can be scaled, multiplied, summed with each other and further modified to achieve a desired effect. For instance, if it is known that a curve behaves differently before and after concentration *θ*, changepoint kernel can be used to provide a reasonable model. If the curve before *θ* can be described by

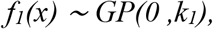

and the curve after *θ* can be described by

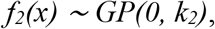

then the desired global GP can be constructed as a weighted average of the two local models

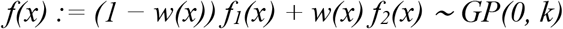

and its kernel function can be composed as

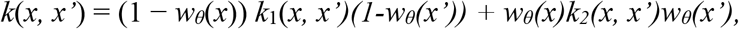

where *w*_*θ*_(*x*) is a function with values between 0 and 1. The further away is *x* from *θ* to the left, the closer is *w*_*θ*_(*x*) to 0 (hence, *k*_1_ will dominate in that region). The further away is *x* from *θ* to the right, the closer is *w*_*θ*_(*x*) to 1, and hence, *k*_2_ will dominate there. Standard choice of the weighting function is the sigmoidal *w*_0_*(x)* = *σ(x)*, providing a smooth transition from *f*_1_ to *f*_2_, centered around zero. However, the weighting function can be modified to shift the transition point from 0 to *θ*: *w*_0_*(x)* = *σ*(*x*) → *w*_*θ*_(*x*) = *σ*(*x* − *θ*), or make the transition more rapid 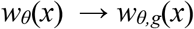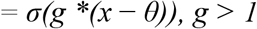 or slower *g* <1. If parameter *g* is a very large number, the transition function can be substituted with a step function *s*_*θ*_(*x*) = 0, if *x*< *θ* and *s*_*θ*_(*x*)=1 otherwise. In practice, if one opts to use the step function as a weight *w*_*θ*_(*x*), it is easier to formulate the changepoint kernel via an if-else statement. In our model formulation, *g* is a parameter and is estimated from data. Dependence of GPs with standard kernels (exponential, squared exponential, Matern, linear) on the values of their parameters are well understood. The novelty of the proposed kernel is in the parameter *g*. Qualitative change of behaviour of a curve, depending on the value of *g*, is presented on Supplement Figure 3. The figure shows how the values of *g* affect the shape of possible curves under such kernel.

In the context of concentration-response curve fitting, the data can be represented as a set of pairs (*x*_*i*_, <u>*y*</u>_*i*_), *i* = 1,…,*N*. Here *x* = *x*_*i*_ is the value of the unique input coordinate (concentration), <u>*y*</u> = <u>*y*</u>_*i*_ is the corresponding mean response, and *N* is the number of unique concentrations (see Figure 2). For PROTACs it is assumed that the expected response is constant at low concentrations up to a threshold concentration *θ* (the concentration threshold at which a chemical induces an effect, i.e. PoD). Linear kernel with no slope is appropriate to model such behavior (*k*_1_ = 0) of the model to the left from *θ*. For concentrations above *θ*, the expected response is assumed to vary smoothly as a function of concentration, and it can both grow or decline. Hence, exponential or squared exponential kernels would make a reasonable choice for concentrations above *θ*. The kernel *k*_2_=*η*^2^(*x*_*i*_ − *θ*)^2^(*x*_*j*_ − *θ*)^2^*exp*(-(*x*_*i*_ −*x*_*j*_)^2^/*ρ*^2^), defining the function to the right side of the threshold, is a product of the Gaussian and squared linear kernels. The Gaussian kernel *η* ^2^*exp*(-(*x*_*i*_ −*x*_*j*_)^2^/*ρ*^2^), provides smoothness, and squared linear kernel (*x*_*i*_ − *θ*)^2^(*x*_*j*_ − *θ*)^2^ allows the range of plausible values to increase together with the distance from the point of departure *θ*.

**Figure 2:**
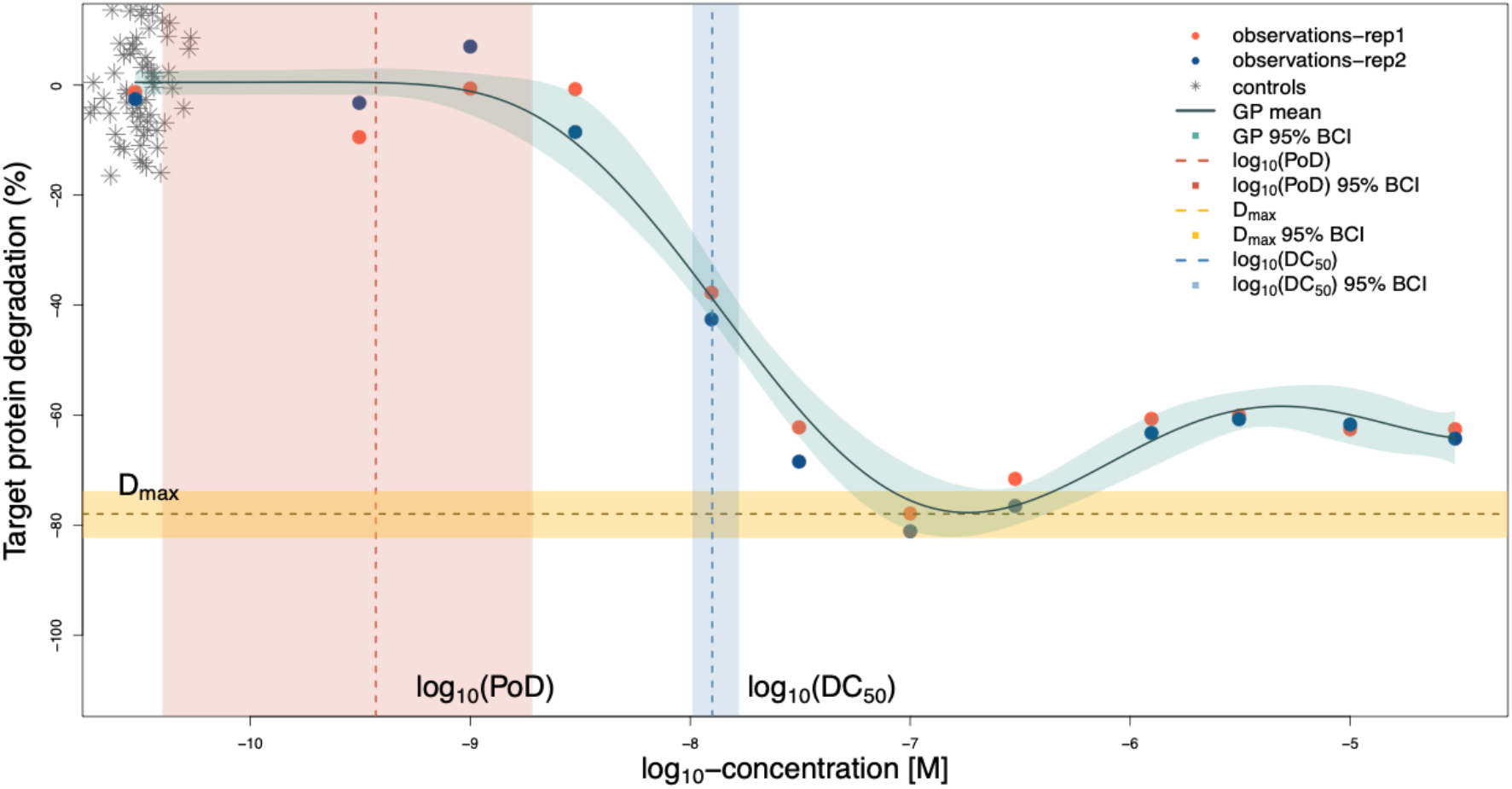
Prediction for one compound: The predicted curve, calculated estimates and uncertainty intervals are shown. The PoD is estimated less precisely compared to the DC_50_ and D_max_. Controls are graphically overlayed at the lowest available concentration for treatments.

As the weighting function we use 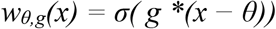, where parameter *g* is inferred from data. Note that the kernel used in^9^ is the special case of the above described kernel, as the weighting function chosen there equals *s*_*θ*_(*x*). By allowing *g* to be a parameter of the model, we allow more flexibility in the curve shapes.

#### GP predictions

Only evaluations of the GP at measured concentrations *x*_*i*_ = *x*_*obs,i*_ are used for inference, but we are interested in finding continuous curves. To calculate curve trajec-tories, we define a grid of points (concentrations) at which we want to make predictions and calculate the values of *f* at the selected locations *x*_*pred,i*_, *i* = 1,…,*N*_*pred*_. Jointly, observed and unobserved values are distributed as a multivariate normal

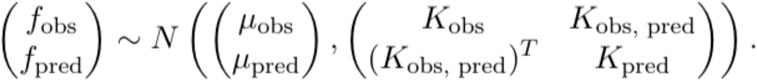

Due to the properties of the multivariate normal distribution, the distribution for the unobserved values given the observed values is given by *N*(<u>*m*</u>, Σ), with <u>*m*</u> = *μ*_*pred*_ + (*K*_*obs, pred*_)^*T*^ * (*K*_*obs*_)−1 * (*f*_*obs*_ − *μ*_*obs*_) and *Σ* = *K*_*pred*_ − (*K*_*obs, pred*_)^*T*^ * (*K*_*obs*_)−1 * *K*_*obs, pred*_. Here *μ*_*pred*_ = *μ* * [1,…,1]^*T*^ is a constant vector, *K*_*obs*_ is the covariance matrix evaluated at observed locations, *K*_*pred*_ is the covariance matrix evaluated at locations for predictions, and *K*_*obs, pred*_ is the covariance matrix evaluated at the pairs of observed and prediction locations.

#### Priors for model parameters

Full Bayesian model formulation requires specification of prior distributions for parameters *μ, σ, σ*_*rep*_, *η, ρ* and *θ*. Baseline response *μ* is assigned the weakly informative prior *μ* ~ *N*(0,0.1). Scale *σ*, governing the concentration-to-concentration variation is assigned the inverse gamma prior *σ* ~ *InverseGamma(1,2),* implying a mode of 1. Scale *σ*_rep_, governing the replicate-to-replicate variation, is assigned the inverse gamma prior *σ*_rep_ ~ *In-verseGamma(1,0.1)* implying a mode of 0.05. In this way we express our belief that replicate-to-replicate variation for the same concentration is smaller than the deviation of measurements from the mean curve. The length-scale, *ρ*, should not be smaller than the minimal distance between concentrations (about 0.5 on the log10-concentration scale), and not larger than the range of concentrations (about 6 on the log10-concentration scale). We use a gamma prior *ρ* ~ *Gamma(50,20)* which does respect the constraints and has a mode of approximately 2.5. This implies that for log10-concentrations between −10.5 and −4.5, correlations are non-negligible at distances of about 40% of the total length of the concentration interval. For the amplitude *η*, we use a normal prior *η* ~ *N(1,1),* and for the threshold *θ*, a uniform distribution between the lowest and highest concentrations *θ* ~ *U*(*x*_*min*_,*x*_*max*_). For degrees of freedom of the Student’s t-distribution *v* we use Gamma(2,0.1). This prior has been proposed in^12^ and is now widely adopted by the Stan community. With mean of 20 and variance of 200 it provides room for a wide range of values for *v*. Our prior for *g is* Gamma(10, 1), which has mean of 10, variance of 10 and hence specifies a plausible range of values as informed by Supplement Figure 3.

#### Different baselines for treatments and controls

Data normalizataion procedure includes the transformation *x*-> (*x*-<*cr*>), where *x* is the raw measured response, and <*cr*> is the median value for controls. As mean and median of controls may not coincide, the difference between them can be non-zero. That is why we keep the parameter *μ* in the model, even if we expect its estimates to be close to zero. In datasets, where a lot of measurements of controls and only few measurements for each treatment dose are available, it can happen that *μ* would represent the mean of controls stronger than the treatment. In cases when observed treatment response at low concentrations is not close to the mean of controls, using *μ* as a baseline (value at low concentrations) for treatments may bias the response curve in the low-concentration area. To avoid such an effect, modification of the model can be used, where the base level of the treatment is estimated by the parameter *μ* and the mean of controls is estimated by a different parameter - *μ*_*c*_. In that case, likelihood of the model takes the form

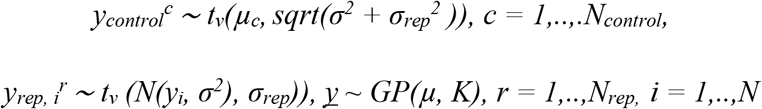

Usually, there won’t be many controls available, and the model with *μ*_*c*_=*μ* would be more appropriate. If this is not the case, however, we suggest using the baseline for treatment different from the mean of controls. For this version of the model we suggest to keep the prior for *μ*_*c*_ as a normal distribution with mean 0 and standard deviation 0.1, and to choose the prior for *μ* as a normal distribution with mean 0 and standard deviation 0.2.

#### Calculating biologically relevant parameters and ranks

In the process of Bayesian inference, we collect samples of the model parameters and the mean GP curve, which form the posterior distribution. If *N*_*iter*_ draws (i.e. MCMC samples) have been saved, each of the curves can be used to derive further statistics, such as biologically meaningful quantities - DC_50_, D_max_ or PoD. We perform the computations based on each drawn GP to create a sampling distribution of the quantities. These distributions reflect uncertainty of the estimates. A PoD is directly represented in the model by its parameter *θ*; D_max_ is calculated as the maximal amount of degradation, i.e. the minimal value on the *y*-axis; and DC_50_ is calculated as the abscissa of the half-maximal degradation point. To find DC_50_ numerically, we first compute its ordinate *y*_DC50_ and then search for a point with the lowest *x*−value on the predictive grid with the minimal difference between the GP and *y*_DC50_.

For each saved MCMC iteration *i* in this process, we create a ranking of compounds based on the magnitude of DC_50_. If (DC_50_)^*i*^_*j*_ is an estimate obtained from iteration *i, i* ∈ (1,*N*_*iter*_) for compound *j, j* ∈ (1, *N*_*cpds*_) (here *N*_*cpds*_ is the total number of compounds), then for a fixed *i*, the *N*_*cpds*_ estimates can be sorted. The order of the sorting yields the ranking of compounds, and the final rank of a compound is its most frequent rank of this compound across all iterations. This procedure is graphically presented on Figure 3. Fitting of the Bayesian model for each compound is taking place independently from other compounds. In the presence of a large number of compounds which need to be compared, this step can be parallelized and will not lead to increased computation time. Once all posterior samples have been collected, the sorting step is applied to each iteration *i* independently and, again, can be parallelized.

**Figure 3:**
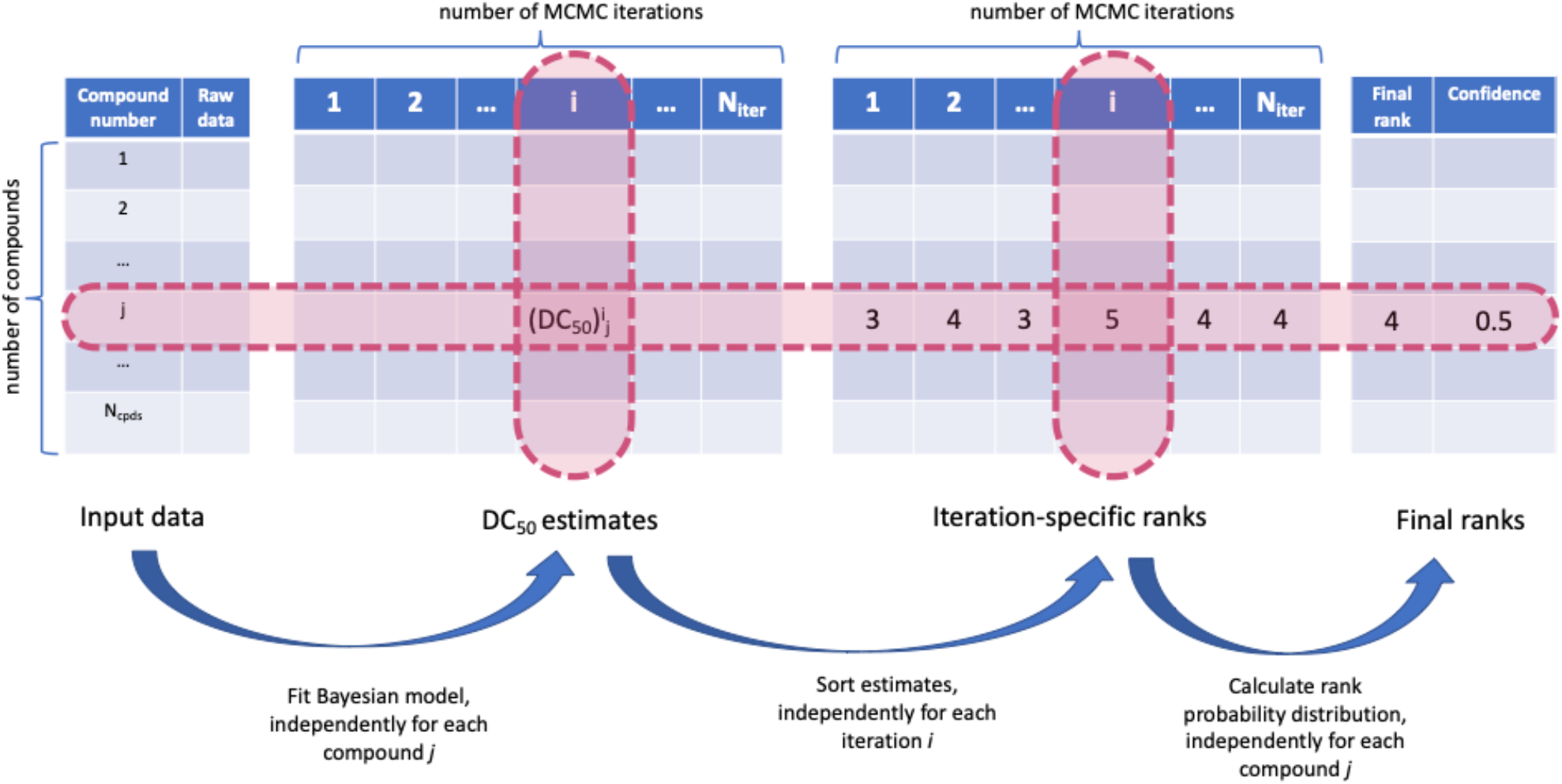
From raw data to compound ranks.

#### Software

The model was fit in Stan^13^ which uses a Hamiltonian Monte Carlo sampler. We used Rstudio^14^ to process the data and interface with Stan. The Bayesplot^15^ library was used to visualize convergence statistics. We used 4 chains of 10000 steps each to produce draws from the posterior. Convergence has been evaluated numerically using *R*-hat and effective sample size statistics. R-hat compares within chain variation and between chain variation and effective sample size is an estimate of the number of independent draws from the posterior distribution. There are no strict thresholds for both statistics, but general guidance is that R-hat should be close to one and effective sample size should be not too small as compared to the total number of samples. Visual inspection of posterior distributions of model parameters (Supplement Figure 4 shows results for one compound) also demonstrated convergence as posterior distributions from different chains agree well. Mean elapsed CPU time (on a laptop with 2.3 GHz Intel Core i5 and 8 GB Memory) for 4 chains was 9.23 seconds with standard deviation of 1.22 seconds. Estimates of model parameters for one compound are presented in Supplement Table 1. Predictions were made on a regular grid of *N*_*pred*_ = 1000 points spaced between the minimal and maximal concentrations. Produced estimates of biologically relevant parameters are presented in Supplement Table 2. The code for fitting the data to one compound using logistic function as a weight function 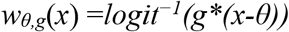 is available on GitHub:

https://github.com/elizavetasemenova/gp_concentration_response.

## Results

First, we fit the model to one compound with 12 concentrations (the dosage currently used in practice) and two replicates and discuss the results in detail. Next, we analyze 9 compounds with 18 concentrations. Data of the two experiments were available, and each experiment contained two replicates. In this way, the combined dataset provides four replicates per concentration for each of the nine compounds. We demonstrate uncertainty in the produced ranking based on DC_50_ and suggest extending this metric to also include D_max_. Finally, we fit the alternative model with different baselines for treatments and controls to all 9 compounds.

### Example of one compound

This dataset contains 12 concentrations with 2 replicates per compound. Figure 2 shows the mean prediction, which closely follows the main trend in the data, and the qunatile-based (95%) Bayesian credible Interval^16^ (BCI; shaded region around the curve). Despite the non-sigmoidal shape, the model accurately detects the lowest part of the curve, which indicates the maximal effect (D_max_). In addition, the model automatically estimates a sensible value for the DC_50_ – similar to where a person might draw a vertical line. The PoD is estimated with less precision but covers the range of concentrations where the curve is largely flat.

### Uncertainty in ranking of compounds

The data in Figure 4 are from 2 experiments at 18 concentrations and with 2 replicates per experiment. This is a very high resolution and would be rarely available in a screening process, and hence they yield tight uncertainty bounds for log_10_(DC_50_). However, even low uncertainty in the estimates translates into the uncertainty of the rankings for many compounds. Figure 5 (a) displays posterior distributions for the log_10_(DC_50_) values and Figure 5 (b) displays the resulting ranks and their uncertainty. Figure 4 shows that for most of the compounds the model provides sensible fits. Compounds 2 and 5 display a similar pattern: there is a discontinuity in the trend of the data. The fitted curve does not exactly fit the points at lower concentrations, or points at about −8.2 concentrations (on the log_10_ scale), but rather finds an average trend between the two extremes. While the model does not overfit, discontinuity in the data is reflected in the uncertainty of the curve estimates in the neighborhood of this concentration. Furthermore, the model behaves robustly with the respect to the outlier of compound 2 (at log_10_-concentration of 6). Compounds ranked as the best (rank=1, 2) and the worst (rank=9, 8) are very distinct, while uncertainty in the ranks is higher for the other compounds.

**Figure 4:**
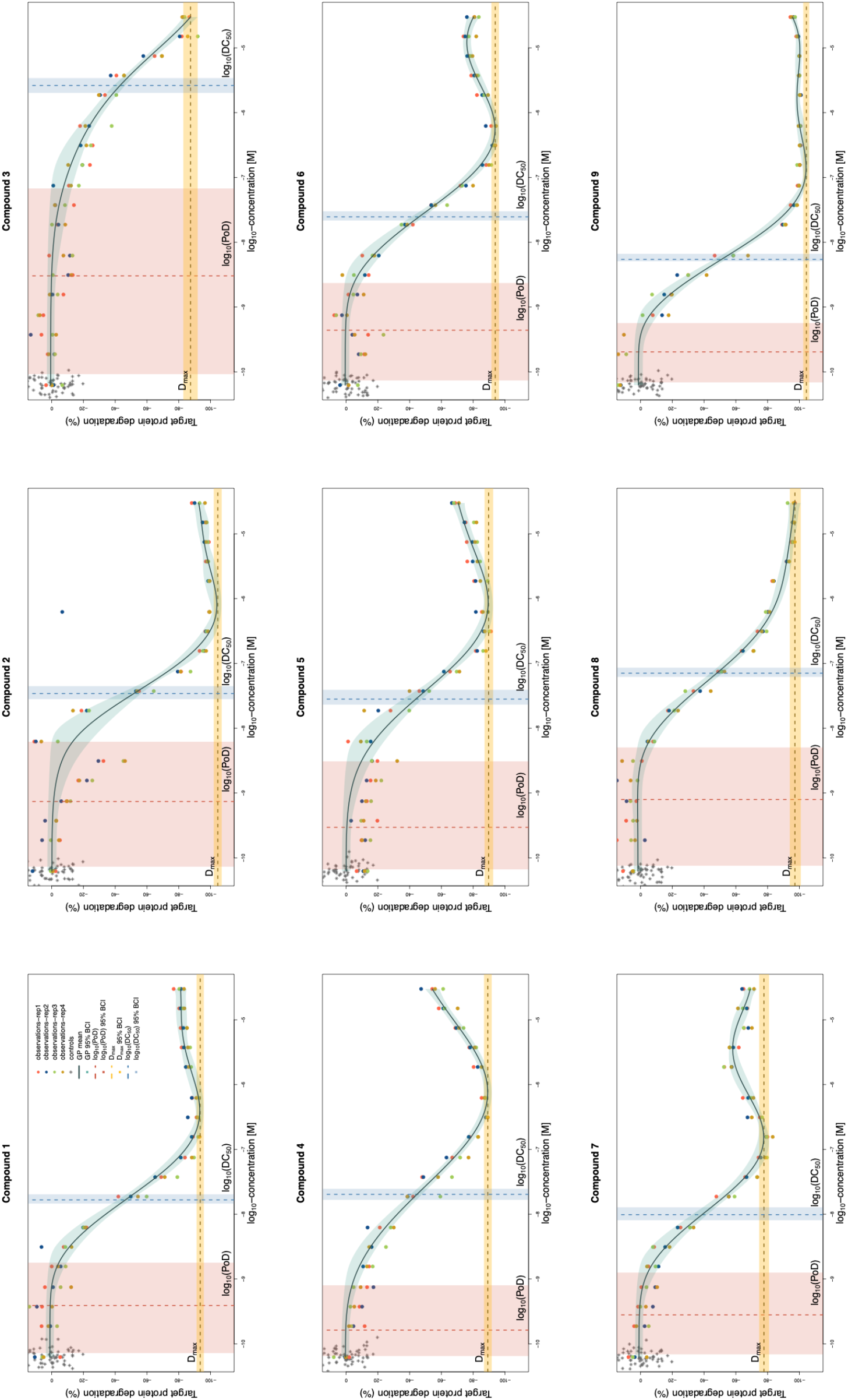
GP model fits of 9 compounds, with 4 replicates each. The flexibility of the model allows it to fit a variety of concentration-response relationships, and despite some non-sigmoidal shapes, the estimated DC_50_ and D_max_ values correspond to what a person might draw by hand. Controls are graphically overlayed at the lowest available concentration for treatments.

**Figure 5 (a):**
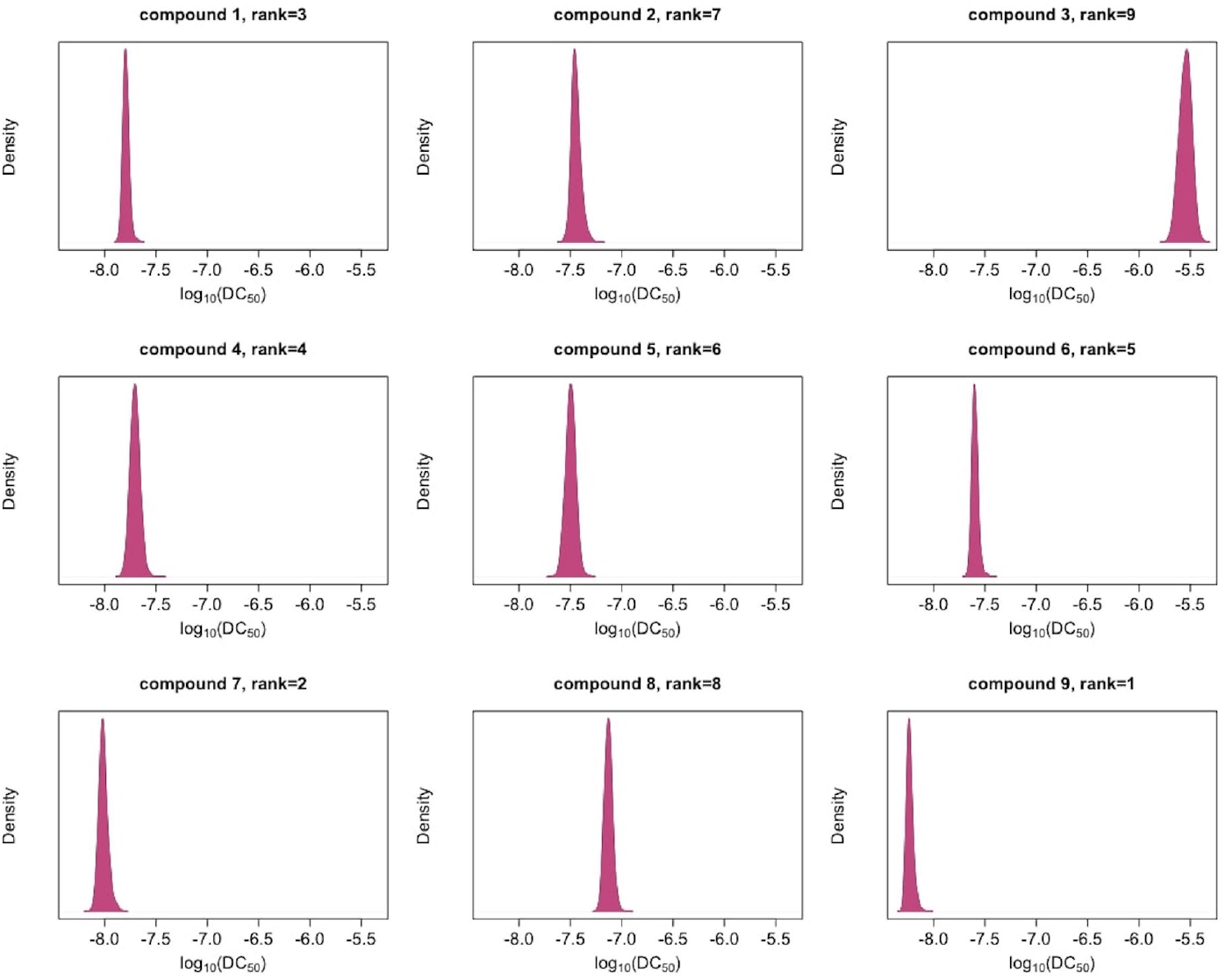
Posterior draws of log_10_(DC_50_) values.

**Figure 5 (b):**
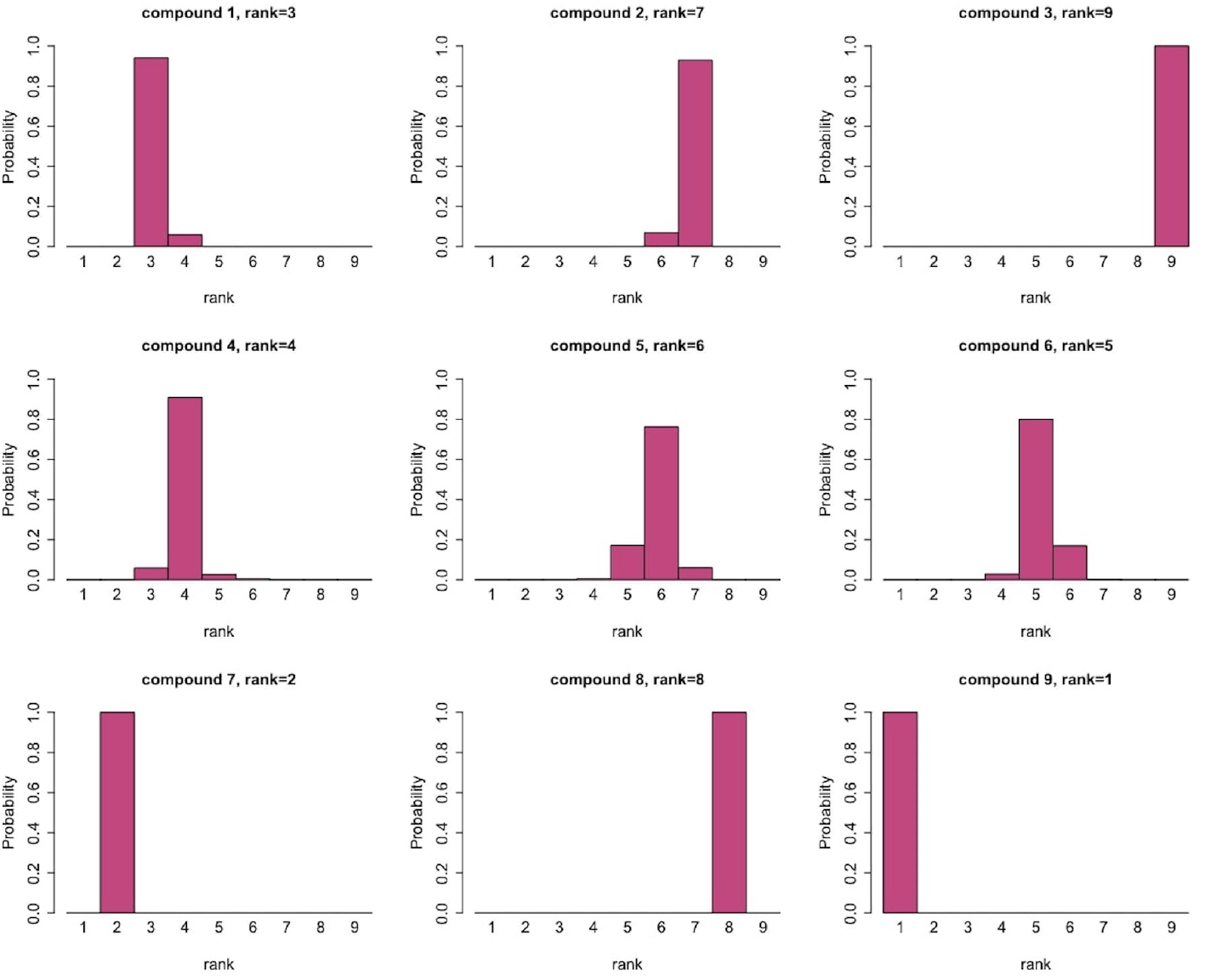
Resulting ranks of compounds together with their uncertainty.

Decisions made solely on DC_50_ might overlook other important properties. Other estimated parameters of the model or the derived biologically meaning quantities can be used for ranking. For instance, the maximal degradation D_max_ carries important information about a compound. There is multitude of ways to choose a ranking measure. Here we present how the combination of DC_50_ and D_max_ can be used. Figure 6 displays the estimates and BCIs for both parameters. Compounds in the bottom left quadrant are preferable. This agrees well with the ranking of compound nine - it has the lowest log10(DC_50_) and log_10_(D_max_); however, compound seven, confidently ranked second according to log10(DC_50_), shows less maximal degradation, and potentially, would be de-prioritized if more information was taken into account. We suggest using a weighted combination of log_10_(DC_50_), and log_10_(D_max_):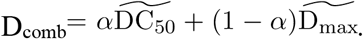. Here 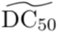 and 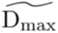 are log_10_(DC_50_) and log_10_(D_max_) re-scaled to the [0,1] range, and *α* ∈ [0,1] is the weighting coefficient, which can be chosen by a scientist to give more preference to DC_50_ (*α* close to 1), weight DC_50_ and D_max_ equally (*α* = 1/2) or to prioritize D_max_ (*α* close to 0). Using the weighted metric, we re-calculate ranks of the 9 compounds according to different values of *α* (Table 1). We have performed the scaling by subtracting the minimal value of all samples and dividing by the difference between minimal and maximal values of all samples.

**Figure 6:**
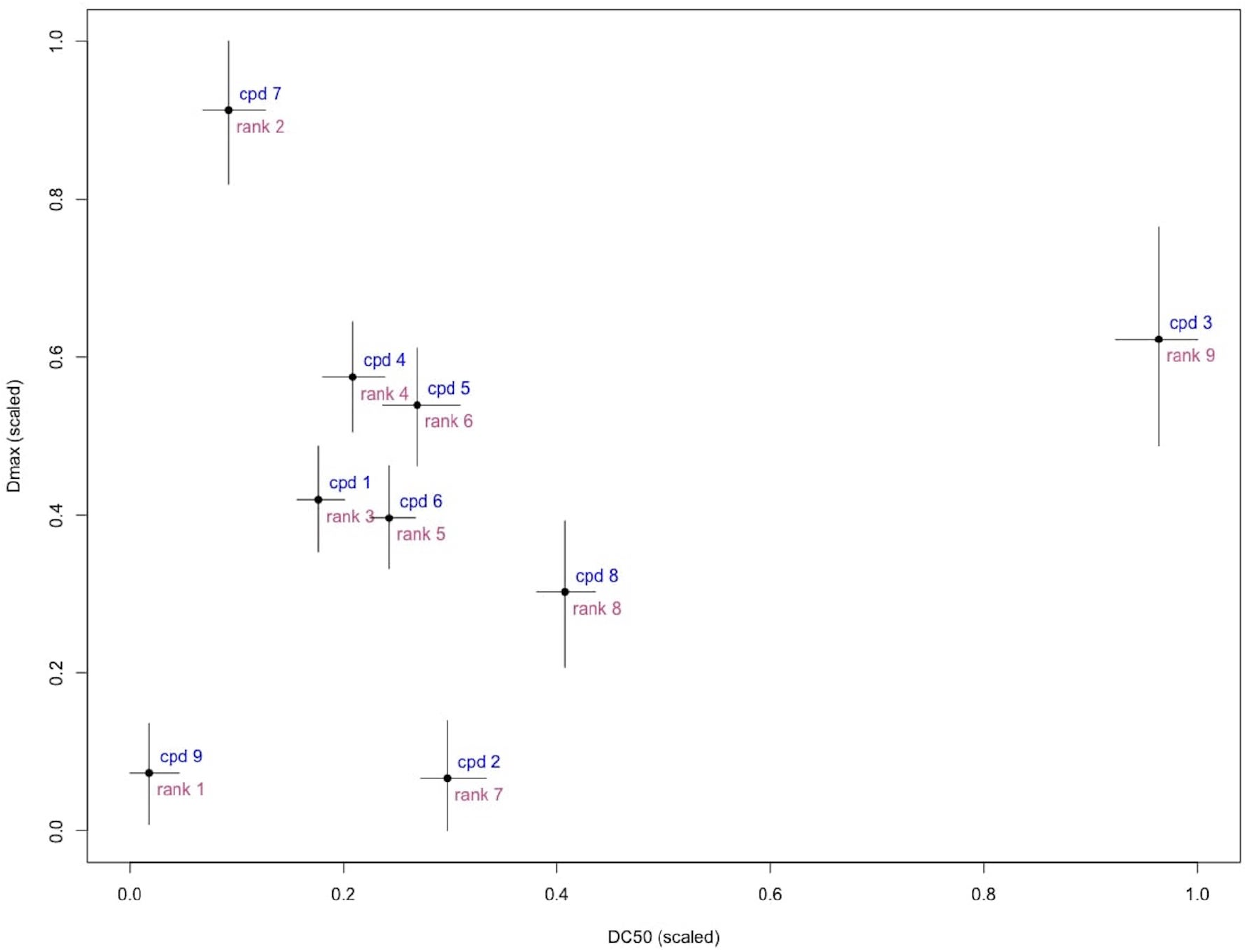
Two or more parameters can be used to rank compounds: Compounds with low values of 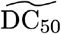 (scaled log_10_(DC_50_)) and 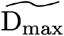 (scaled D_max_) are desirable as they indicate high potency and large effects. Compound 9 is distinguished as the best by these criteria. Error bars are 95% BCI.

**Table 1.**

| $\alpha$ | Compound number | | | | | | | | |
| --- | --- | --- | --- | --- | --- | --- | --- | --- | --- |
|  | 1 | 2 | 3 | 4 | 5 | 6 | 7 | 8 | 9 |
| Compound's rank (and its relative frequency) |  |  |  |  |  |  |  |  |  |
| 1 | 3 (0.97) | 7 (0.94) | 9 (1.00) | 4 (0.95) | 6 (0.88) | 5 (0.91) | 2 (1.00) | 8 (1.00) | 1 (1.00) |
| 2/3 | 3 (0.83) | 2 (0.84) | 9 (1.00) | 5 (0.78) | 7 (0.48) | 4 (0.92) | 6 (0.50) | 8 (0.70) | 1 (1.00) |
| 1/2 | 3 (0.89) | 2 (1.00) | 9 (1.00) | 6 (0.63) | 7 (0.77) | 4 (0.81) | 8 (0.99) | 5 (0.68) | 1 (1.00) |
| 1/3 | 3 (0.47) | 2 (1.00) | 9 (0.99) | 6 (0.59) | 7 (0.59) | 5 (0.42) | 8 (0.99) | 5 (0.37) | 1 (1.00) |
| 0 | 5 (0.63) | 1 (0.62) | 8 (0.70) | 7 (0.52) | 6 (0.60) | 4 (0.61) | 9 (1.00) | 3 (0.94) | 2 (0.62) |

Ranks of 9 compounds and uncertainty based on the combination metric
$\alpha \widetilde{DC}_{50} + (1 - \alpha) \widetilde{D}_{max}$ for different values of $\alpha$ .

All nine compounds presented in Figure 4 are active compounds, i.e. they respond to the increasing concentration in a significant way. We have tested our model on simulated “flat” data to verify that it would deal with inactive compounds appropriately. For this, we have simulated responses with a mean equal to the mean of controls and a standard deviation which equals 1.1 standard deviation of controls. Supplement Figure 5 displays the fits of the Bayesian Hill’s model and the changepoint GP model to such data. Changepoint GP produces a nearly flat fit with D_max_ estimates close to the mean of the simulated data. Results of the model with different baselines for treatments and controls are presented on Supplement Figure 6. Uncertainty in *μ*_*c*_ is higher than in the model with equal baselines both due to a wider prior for *μ*, but also due to less data available for its estimation.

## Discussion

We have used a changepoint GP concentration-response model, coupled with Bayesian inference of its parameters to flexibly fit PROTAC CR curves. A Bayesian approach is advantageous for small amounts of data and can represent uncertainty of the parameters and derived estimates in the form of distributions. A GP model does not require strong assumptions on curve shapes, as parametric models do. The model is stable to outliers due to using a Student’s distribution. The presence of outliers can be diagnosed using the estimates of the parameter *v*. Non-identifiability is a frequent problem for GPs: often only one of the length-scale or amplitude parameters can be identified, while the other parameter needs to be fixed for the analysis. We have used an informative prior for length-scale and have not encountered problems under the specified settings.

We have demonstrated how uncertainty in the estimates of DC_50_ influences uncertainty in ranks and have proposed a single metric to capture both DC_50_ and D_max_. This metric involves a weighting parameter that allows a scientist to choose their decision-making approach and tackle ranking in a quantified manner. We have demonstrated that the ranks of the worst (rank=9) and the best (rank=1) compounds are fairly stable with respect to the varying weighting coefficient, while ranks of less clear leaders (e.g., compounds number 2 and 7) may change their ordering. Currently, decisions are mainly made based on DC_50_, which corresponds to *α* = 1. The weighting can be extended to more parameters, e.g. DC_50_, D_max_, and *θ* if they all represent interest for screening; the combination metric would weight all three estimates 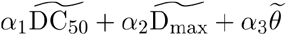 with *α*_1_ + *α*_2_ + *α*_3_ = 1, *α*_*k*_ ∈ (0,1). For parametric models, such as a sigmoidal or double-sigmoidal, the area under curve (AUC) can be used as a single measure of compound’s potency. This approach, however, is less applicable to GP fits since such curves might experience local fluctuations. These fluctuations would have little influence on the estimates of parameters such as DC_50_, but cumulatively might contribute non- negligible amounts of AUC leading to erroneous conclusions. A potential solution could be computing AUC for GPs only in the domain between the minimal concentration and the first concentration where the maximal effect D_max_ is achieved. In our example we have mapped DC_50_ and D_max_ by scaling. Alternative approaches can be used, such as desirability functions^17^.

The proposed model has been used internally to answer a set of experimental design questions, such as “Is it better to have more concentrations, or more replicates?” A series of comparisons has been done between 4,3,2 and 1 replicate for 18 concentrations, and between different number of concentrations (9 and 18) for the same number of replicates. An example of 18 and 9 concentrations (2 replicates each), for the same compound as Figure 2 is shown in the Supplement Figure 2. The plot shows slightly more uncertainty in the model fit produced using 9 concentrations, while the general shape of the fit and DC_50_ estimate remain close. Model fits based on 4,3, and 2 replicates for the same compound are presented on Supplement Figure 8. Both the curve shapes and amount of uncertainty are similar for all three fits. Our conclusion from this exercise, evaluating the trade-off between number of replicates and number of concentrations, was that more concentrations is preferred to more replicates and that 12 concentrations with 1 replicate is sufficient to produce a robust ranking.

The changepoint kernel, used in this work, fits a flat curve to the data to the left side of the PoD and does not use the mechanistic understanding of the model otherwise. As a result of that, some GP fits might display minor local fluctuations at higher concentrations, which a parametric model would not do. Nevertheless, these fluctuations would not have major impact on the estimates of the estimated parameters and ranking. If it is important to capture more mechanics in the curve-fitting step, it can be done by further expansion of the changepoint kernel. For instance, in case of PROTACs, one would expect the curve to flatten out after the hook. The changepoint kernel, in this case, would have three GP components (flat linear, smooth, and flat linear) and two inflexion point *θ*_1_ and *θ*_2_. Derivations of the kernel in this case, are provided in the Supplement. Another solution to the local fluctuations at high concentrations could be provided by a concentration-dependent length-scale, such as, for instance *ρ*^2^(1+(*x*_*i*_ −*θ*)^2^)(1+(*x*_*j*_ −*θ*)^2^)^18^.

GP models are computationally more involved than parametric models. Running Markov Chain Monte Carlo samplers might take longer than applying traditional (frequentist) parametric methods. As a result, incorporating Bayesian GP models into workflows might be less straightforward. However, as computational tools evolve, this limitation will be gradually diminished. Furthermore, frequentist optimization techniques have their own difficulties, such as subjective choice of initial values, and might have convergence issues.

The most time-consuming steps in the compound ranking process is fitting of the Bayesian model and predicting the amount of degradation at unobserved concentrations. As the model can be fit to each of the compounds independently, it takes the same amount of time to fit all compounds, as it takes to fit one compound. Fitting of one compound on our machine takes, on average, 9 seconds (with standard deviation of 1.2 seconds between 4 chains). As the model is fit to the data for each compound independently, the process can be parallelized without additional time cost. The sorting procedure that provides the ranks is not specific to Bayesian methods and would be needed in a standard ranking procedure as well. The overall computation time is a trade-off with the number of iterations for Bayesian model fitting and the number of concentrations at which the response needs to be predicted.

Among the nine presented compounds, one of them (compound 3) could not have reached its D_max_. The estimates produced by the changepoint GP model reflect the uncertainty in a wider BCI as compared to other compounds. Fit of compound 3 does not have an asymptote at higher concentrations as values at these concentrations keep declining monotonically. This is an issue of the range of observed concentrations, and not of the model. Width of the uncertainty interval for D_max_ here is wider than for other presented compounds fitted by the changepoint GP model. For comparison, we have also produced Bayesian fits of the 4PL (4-parameter) model to the data, accounting for the same sources variation as described above, and the mean curve modelled as *y* = *d* + (*a-d*)/(*1 + exp(-b(x-c)))*, where *d* denotes the amount of degradation at zero concentration, *a* denotes the D_max_, *c* stands for log_10_(DC_50_), x is log_10_-dose and b is the Hill’s slope. More details and produced fits for all 9 compounds are available in the Supplement Figure 9. For compound 3, the BCI of D_max_ produced by the 4PL model is (−90.11, −85.93) and has the width 4.18; BCI produced by the changepoint GP model is (−91.66, −82.49) and has the width 9.17; non-Bayesian fit of the 4PL model (obtained using the *drc* R-package^19^) produces a CI of (−309.2, −67.86) of width 242. While the width of the BCI by the changepoint model might be more narrow than intuitively expected, the CI by the frequentist model is too wide: its upper limit of approximately −67 is much higher than the observed degradation at high concentrations (below −80). Additional quantities can be calculated to resolve this issue and verify whether D_max_ has been attained. The derivative of the function, approaching D_max_ should be nearing zero, and hence the absolute value of the slope of the curve should be small. The magnitude of this derivative can be incorporated into the estimation of uncertainty of D_max_. In this work we have used data, pre-processed by GeneData via the normalization step. The shift *x*-<*cr*> moves raw data *x* up or down by the mean of the controls to make all curves start at the value of approximately *0*. The distribution of neutral controls is tight, and its mean and median are not far from each other. The estimates of parameter *μ* confirm that: e.g. Table 1 in the Supplement shows that this is a very small number and its credible interval includes *0*. The entire transformation (*x*-<*cr*>)/(<*sr*>-<*cr*>) ensures that all data points lie approximately between *0* and *−100*. Hence, normalization affects curve-to-curve range, and enables comparability of curves and produced estimates. For the case when treatment response at low concentrations is not close to the median of controls, we have suggested an alternative model with different baselines of treatments and controls. For compounds with lower base level *μ* than in model with equal baselines, e.g. compounds 2 and 5, POD has shifted to the right; on the contrary, POD of compounds with higher *μ,* e.g. compounds 8,9, POD has shifted to the left.

Unlike parametric models, it is more difficult to link GPs to a mechanistic understanding. And sometimes models might display minor local fluctuations, which a parametric model would not have. Nevertheless, these fluctuations would not have major impact on the parameters estimates and ranking. We believe that this approach has a lot of potential and can be used by pharmaceutical companies by taking steps towards its automation. If the assumption about possible curve shapes holds, important choices to make are the priors, and degrees of freedom in the Student’s t-distribution.

## Acknowledgments

Elizaveta Semenova is a fellow of the AstraZeneca postdoctoral program. We thank Joe Reynolds for sharing their code from^9^. We thank the two anonymous reviewers whose comments/suggestions helped improve and clarify this manuscript.

## Supplement

**Table 1:** Summary of the fitted model for one compound.

|  | mean | sd | 10% | 50% | 90% | n_eff | Rhat |
| --- | --- | --- | --- | --- | --- | --- | --- |
| $\sigma_{\text{rep}}$ | 0.11 | 0.02 | 0.08 | 0.11 | 0.14 | 1767.15 | 1.00 |
| $\sigma$ | 0.29 | 0.03 | 0.25 | 0.29 | 0.33 | 3494.05 | 1.00 |
| $\mu$ | 0.02 | 0.04 | -0.03 | 0.02 | 0.06 | 4795.64 | 1.00 |
| $\rho$ | 2.55 | 0.34 | 2.14 | 2.53 | 2.99 | 4073.17 | 1.00 |
| $\eta$ | 0.63 | 0.43 | 0.20 | 0.52 | 1.21 | 2538.65 | 1.00 |
| $\nu$ | 26.66 | 14.47 | 11.20 | 23.66 | 44.80 | 4361.59 | 1.00 |
| $g$ | 9.89 | 3.19 | 6.12 | 9.51 | 14.14 | 5110.81 | 1.00 |

**Table 2:**
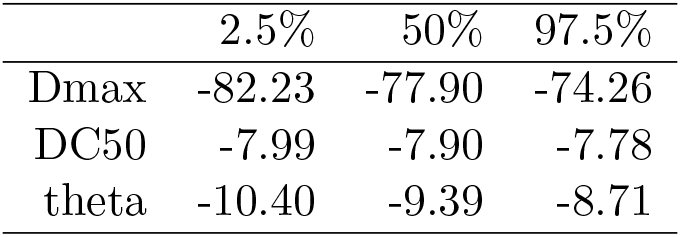
Summary of the estimated biologically relevant model parameters for one compound.

**Figure 1:**
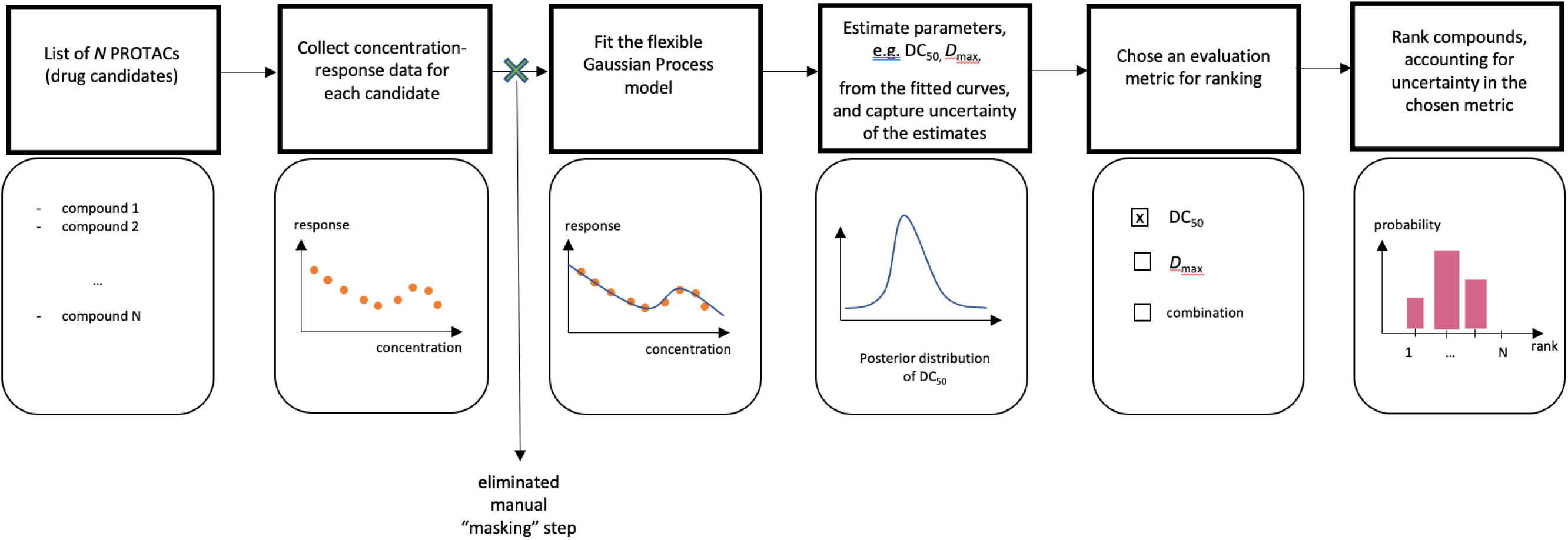
The process of compound ranking.

**Figure 2:**
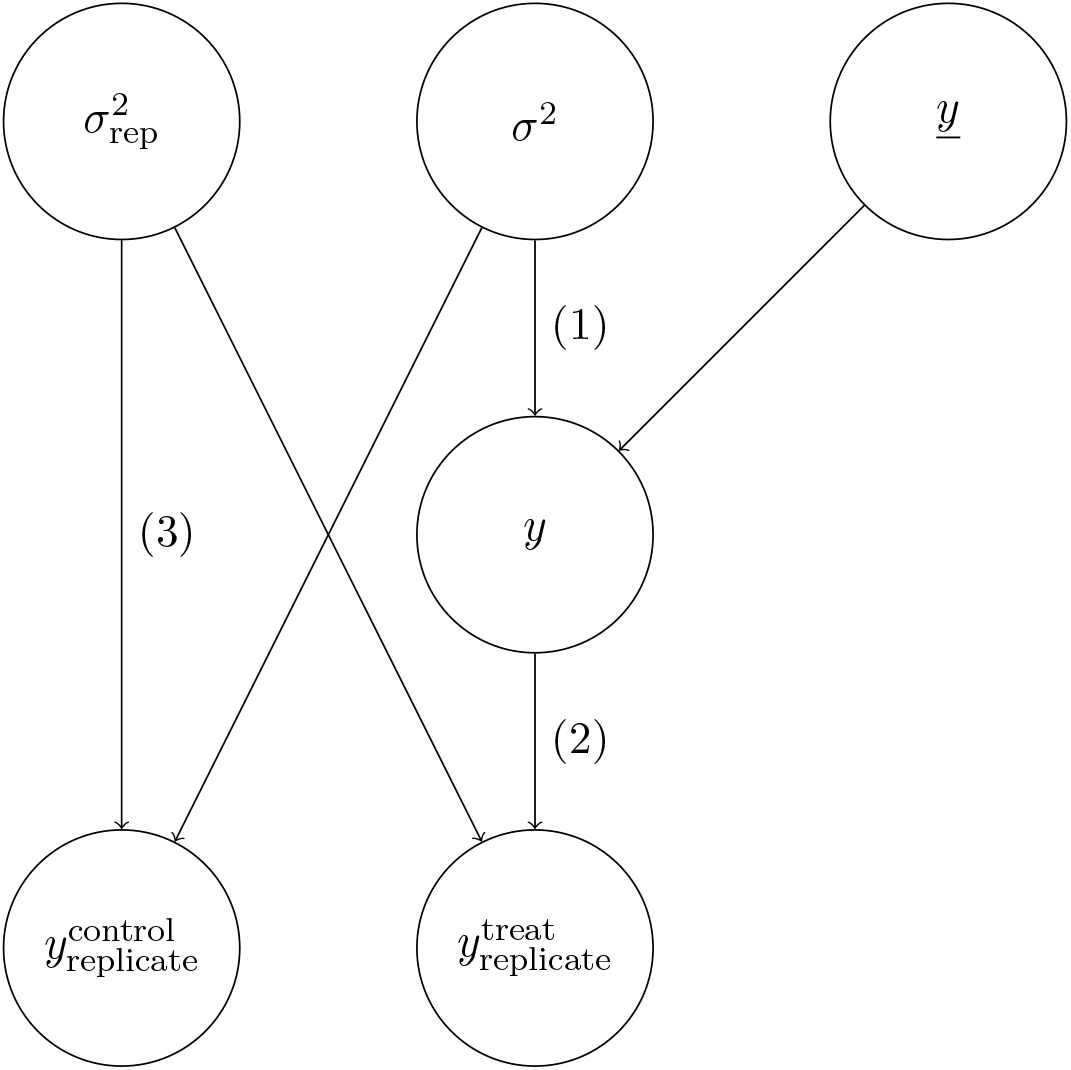
Sources of variation for treatment and control data: <u>*y*</u>-mean predicted curve, *σ* - curve uncertainty, *σ*_rep_ - variation between replicates, and

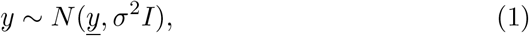

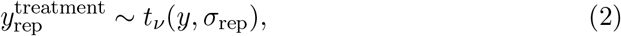

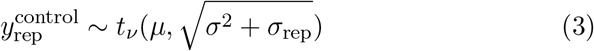

**Figure 3:**
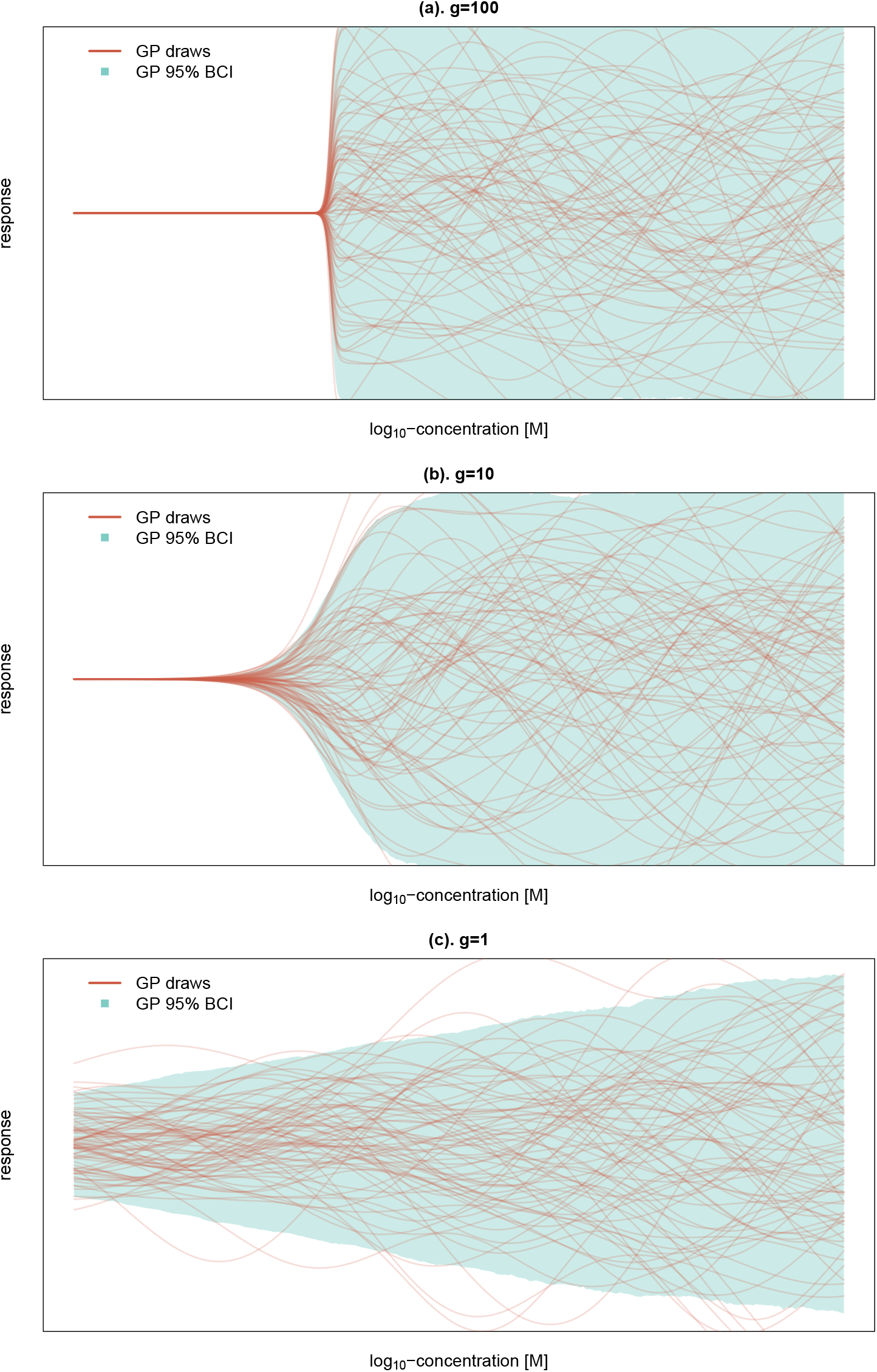
Example realisations of a changepoint Gaussian process - qualitative change in behavoiur as dependence on the parameter *g*: simulations from changepoint kernel model, corresponding to the family of weighting functions 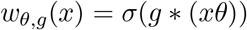.

**Figure 4:**
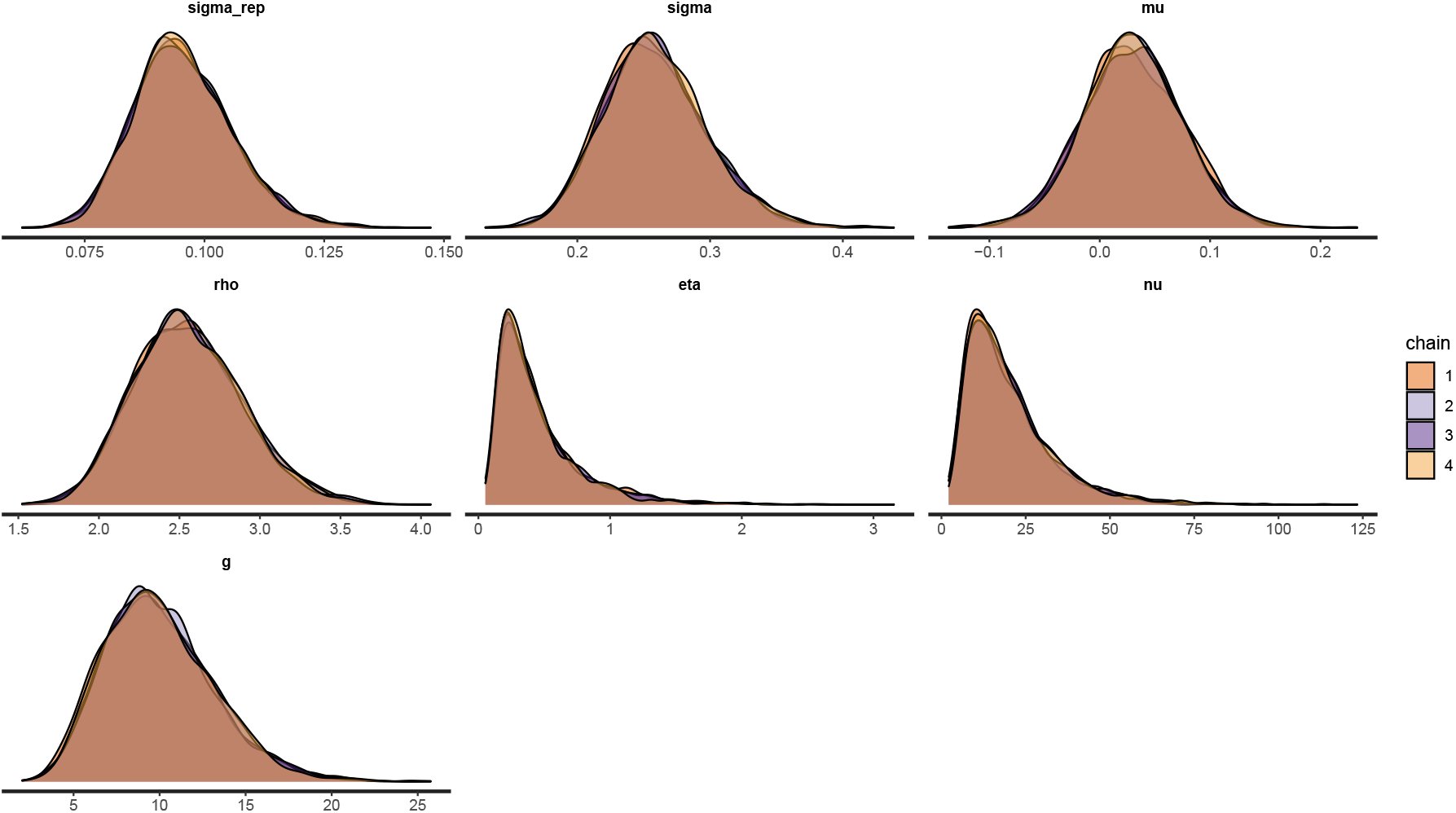
Posterior distributions of model parameters from different chains agree very well and show convergence.

**Figure 5:**
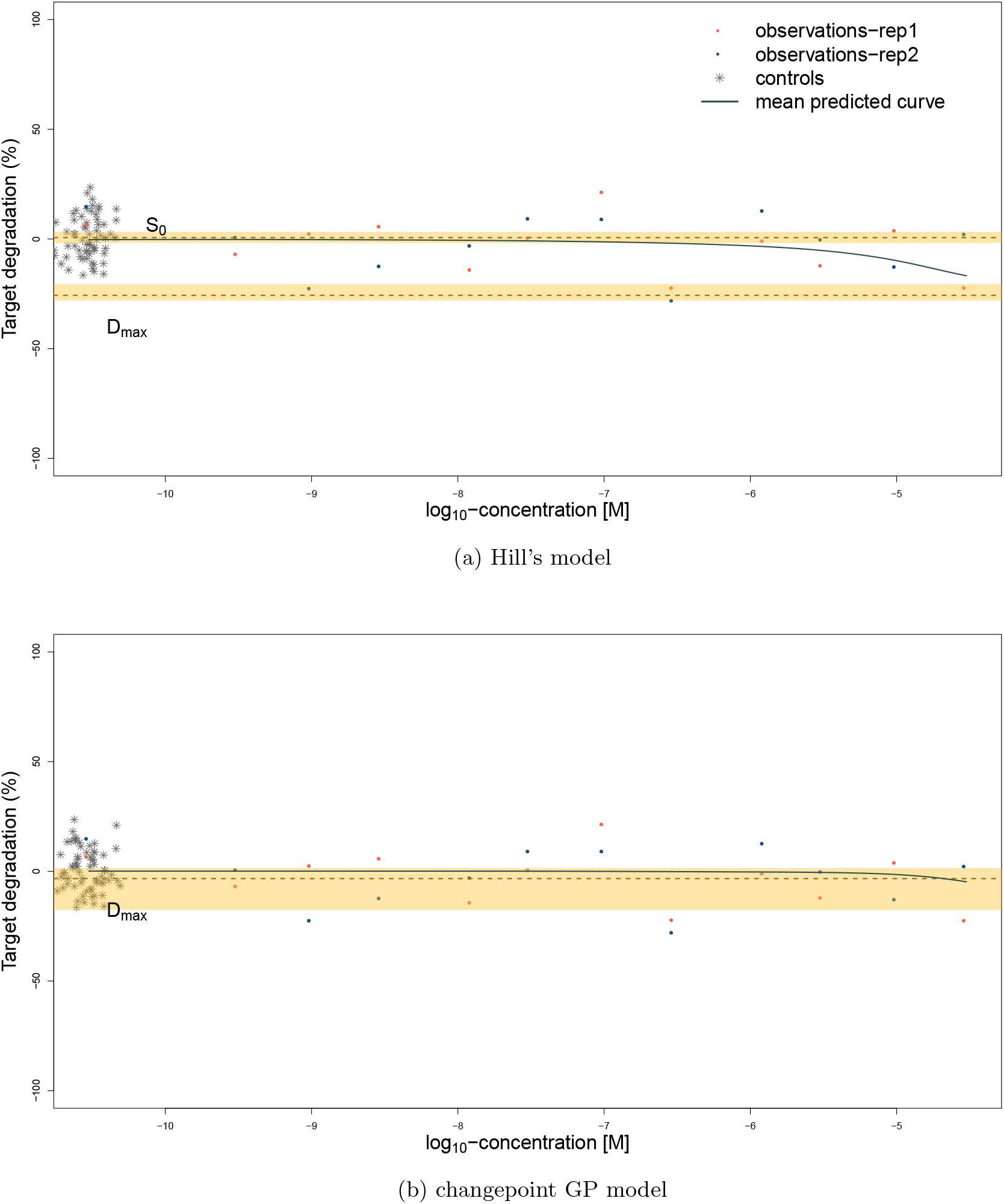
Simulated ‘flat’ response fitted with two models: (a). Hill’s (4-parameter 4PL) model, (b). changepoint GP model.

**Figure 6:**
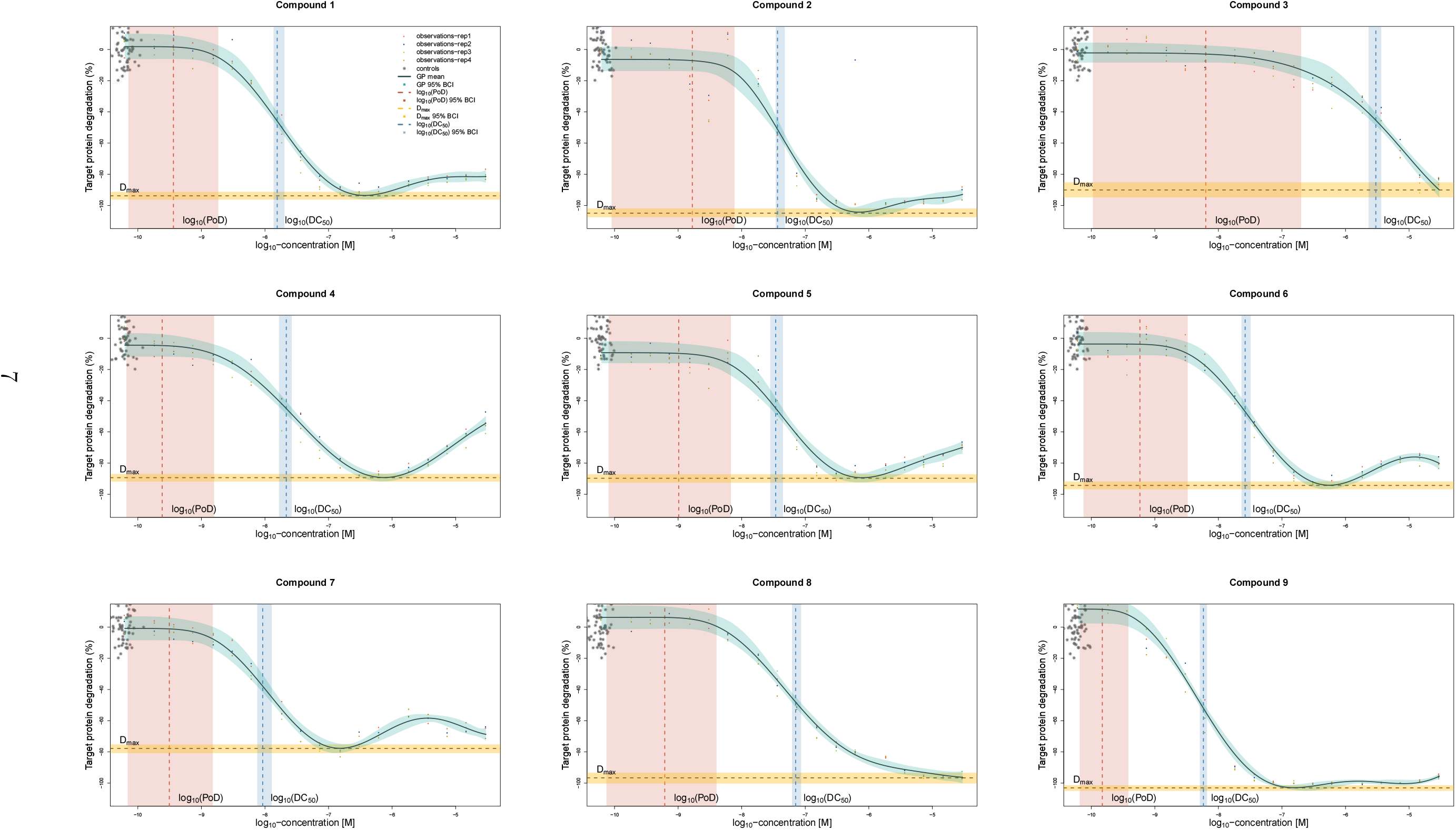
Fits of the changepoint GP model with different means for treatment and control.

**Figure 7:**
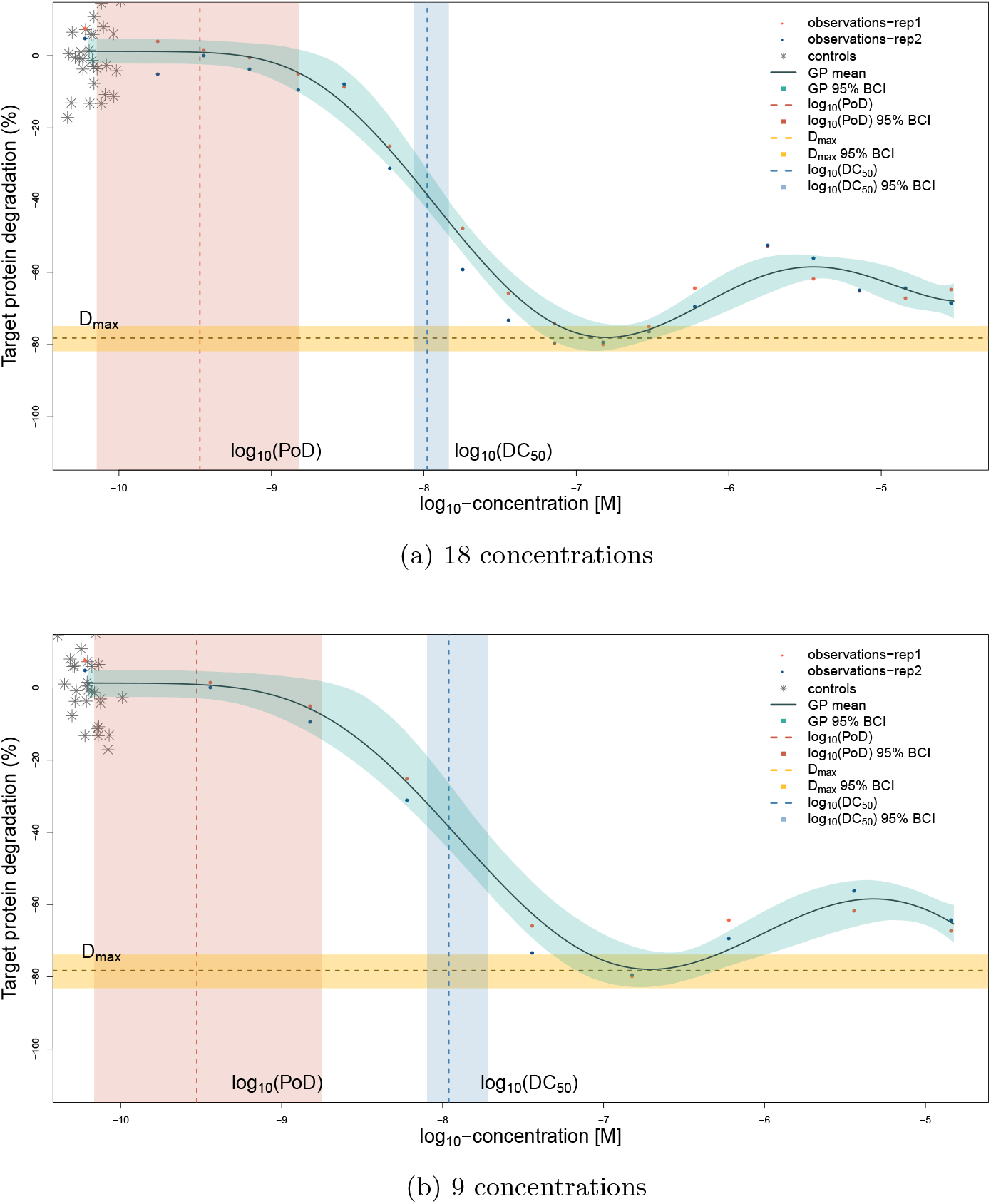
Example of one compound: fitted model and calculated estimates based on 18 and 9 measured concentrations in 2 replicates each.

**Figure 8:**
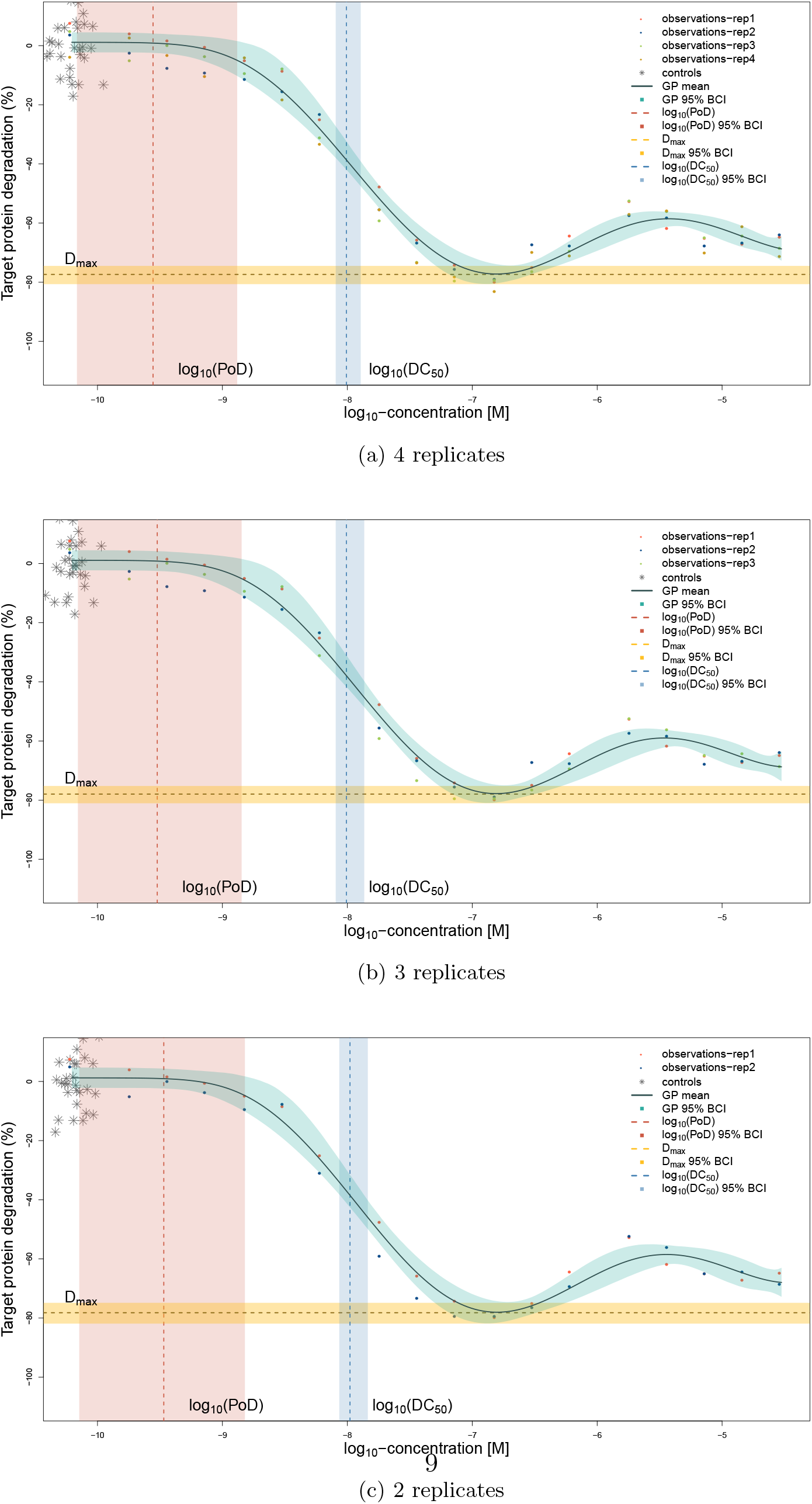
Example of one compound: fitted model and calculated estimates based on 4,3 and 2 replicates, with 18 concentrations each.

**Figure 9:**
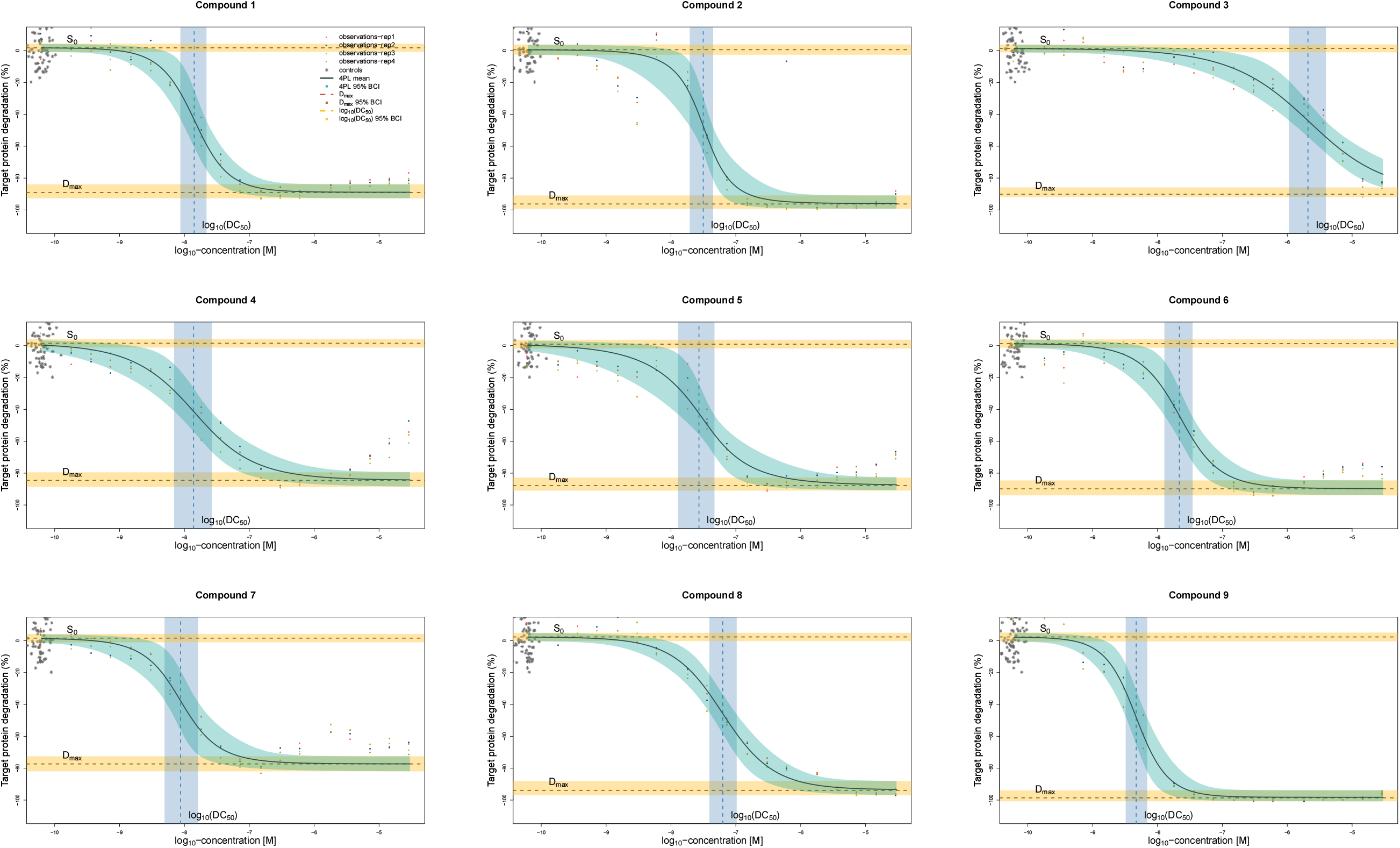
4PL model fits of 9 compounds, with 4 replicates each.

## Model Summary

### Priors for the changepoint Guassian Process model

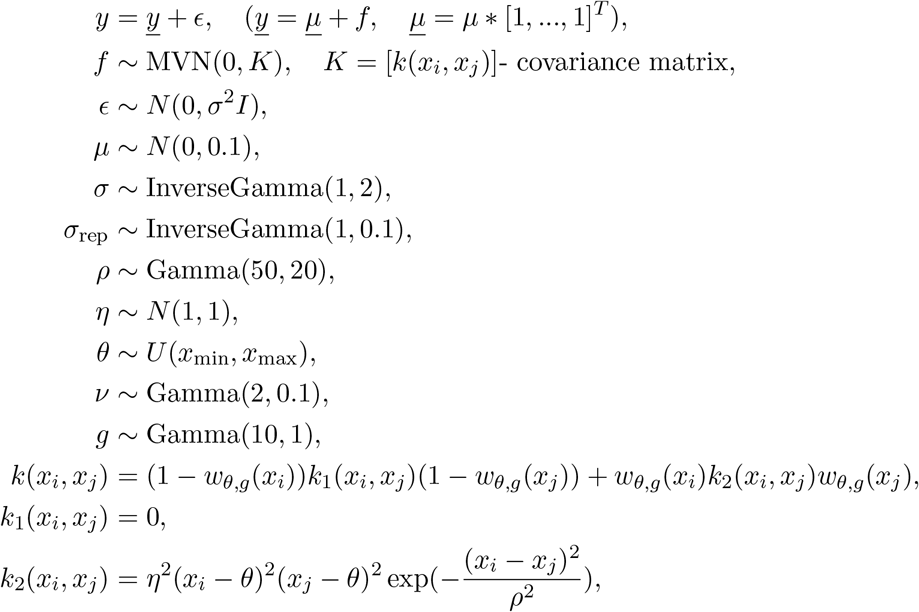

where 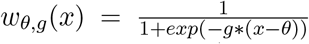 is the weighting function. Note that if 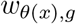 is a step function (i.e. *g* = ∞) at *x* = *θ*, the special case of the kernel becomes

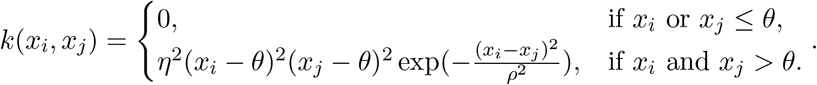

### Likelihood of the changepoint Guassian Process model

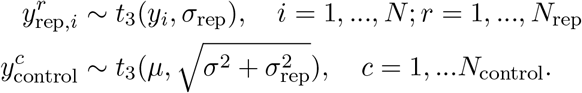

### 4PL (Hill’s) model

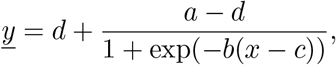

where *d*- degradation at zero concentration, *a* = *D*_max_, *c* = log_10_(DC_50_), *x* - log_10_(dose), *b* - Hill’s slope. Priors, selected to produce fits on Supplement Figure 4 are *σ* ~ InvGamma(1, 2), *σ*_rep_ ~ InvGamma(1, 0.1), *d* ~ *N* (mean(*y*_control_), 0.1), *a* ~ *N* (mean(*y*_treatment_), 0.1), *b* ~ *N* (1, 5).

### Changepoint models

#### Two functions

Changepoint between two GPs and one transition point:

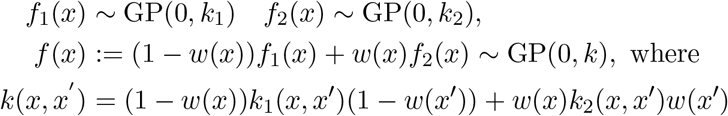

These formulas are easy to understand if we chose the transition function as a step function *w*(*x*) = *s*_*θ*_(*d*), where *d* = *x* − *θ* and

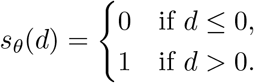

The most popular choice of *w*(*x*) is the inverse logit (sigmoidal) function. A steep sigmoidal function *σ*(*g* * *x*), *g* > 1 is a good approximator of the step function.

#### Three functions

Changepoint between three GPs with two transition points *θ*_1_*, θ*_2_ can be constructed in a similar manner: we need to construct a function which behaves as *f*_1_(*x*) for *x* ≤ *θ*_1_, as *f*_2_(*x*) for *θ*_1_ < *x* ≤ *θ*_2_ and as *f*_3_(*x*) for *x* > *θ*_2_. This can be achieved by

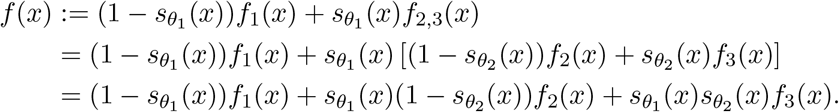

The kernel of this representation can be derived as

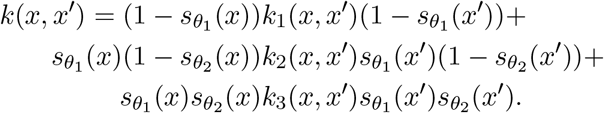

To enforce the hook behaviour with flat curve at low and high concentrations and flexible shape in-between, a reasonable choice of kernels would be linear (without slope) choices for *k*_1_ and *k*_2_ and a smooth kernel for *k*_2_.

